# Mechanism-Aware Inductive Bias Enhances Generalization in Protein-Protein Interaction Prediction

**DOI:** 10.1101/2025.07.04.663157

**Authors:** Shuchen Deng, Xuanjun Wan, Zichun Mu, Sheng-You Huang, Chengfei Yan

## Abstract

Robust prediction of protein–protein interactions (PPIs) requires models that generalize beyond the training distribution. Here, we present PLMDA-PPI, a mechanism-aware framework that incorporates biophysical inductive biases into a dual-attention architecture with residue-level contact supervision applied to protein language model embedded geometric representations. This design enables the model to jointly learn global interaction likelihoods and the residue pairs mediating interactions, producing interpretable, mechanism-grounded predictions. PLMDA-PPI demonstrates strong out-of-distribution generalization, substantially outperforming lightweight deep learning models (D-SCRIPT, Topsy-Turvy, TT3D, and TUnA) and achieving comparable or even superior performance to computationally intensive methods (AF2Complex, RF2-Lite, and RF2-PPI) derived from AlphaFold2 and RoseTTAFold2, while requiring orders of magnitude fewer computational resources. These results show that incorporating mechanistic priors as architectural inductive biases enhances generalization, interpretability, and computational efficiency, providing a principled foundation for AI-driven prediction of complex biological interactions.

## Introduction

Protein–protein interactions (PPIs) play essential roles in nearly all biological processes, underpinning cellular function, disease mechanisms, and therapeutic target discovery^1–3^. Identifying these interactions is critical for understanding molecular mechanisms and enabling drug development. However, experimental mapping of PPIs remains labor-intensive and incomplete, even for well-studied organisms such as human, where validated interactions cover only a small fraction of the complete interactome^4–7^. High-throughput screening approaches, while extensive, often suffer from high false-positive rates, further limiting data quality^8, 9^. Consequently, computational prediction of PPIs has become an essential tool for systematically characterizing interactomes.

Recent advances in deep learning have enabled promising results in PPI prediction^10–12^. Nevertheless, existing approaches face critical limitations. Training these models requires large amounts of high-quality interaction data; however, most datasets are derived from literature-mined PPI databases, such as STRING^13^, which are biased toward well-studied proteins and contain many false positives. As a result, both training and evaluation can exhibit systematic biases^14–16^. For example, Bernett et al.^14^ showed that when sequence similarity between training and test sets is minimized, nearly all previously reported high-performing PPI prediction models lose predictive power. Further analysis revealed that apparent high performance often resulted from information leakage: random splitting of PPI datasets allowed overlapping proteins or proteins with high sequence similarity to appear in both sets. Consequently, many models relied on shortcuts, such as sequence similarity or node degree in the PPI network, rather than learning truly generalizable features. These models therefore fail to generalize effectively to novel proteins with low sequence similarity to those seen during training, highlighting the need for approaches that capture mechanistic determinants of PPIs.

Mechanistically, PPIs are driven by physicochemical interactions between interfacial residues, which stabilize complexes in their native conformations to carry out their functions. During protein–protein association, functional PPIs would form distinctive residue-level contact signatures that are absent in non-functional protein pairs. We hypothesize that a PPI prediction model capable of capturing these residue-level interaction patterns inherently learns the underlying biophysical principles of protein association, thereby improving its ability to generalize to unseen proteins.

Motivated by this insight, we developed PLMDA-PPI, a mechanism-aware framework for PPI prediction. The model integrates biophysical inductive biases via residue-level contact supervision into a dual-attention architecture^17, 18^. Given a protein pair, PLMDA-PPI first predicts an inter-protein residue–residue contact map using protein language model–embedded geometric graphs^19^, which is then fused with sequence embeddings to predict global interaction likelihoods. Both modules are trained jointly, allowing the model to capture key interfacial residue pairs and support interpretable, mechanism-grounded PPI predictions We trained PLMDA-PPI on structure-informed PPIs from the Protein Data Bank (PDB)^20^ and evaluated it on multi-species PPIs from the HINT database^21^. Fine-tuning on human PPIs from HINT further improved predictive performance while maintaining robustness under strict sequence-dissimilarity constraints. Across benchmarks, PLMDA-PPI consistently outperforms recent lightweight models (D-SCRIPT^22^, Topsy-Turvy^23^, TT3D^24^, and TUnA^25^) and achieves comparable or even superior performance to computationally intensive methods (AF2Complex^26, 27^, RF2-Lite^28^, and RF2-PPI^29^) derived from AlphaFold2^30, 31^ and RoseTTAFold2^32^, while requiring substantially fewer computational resources. These results demonstrate that incorporating mechanistic priors as architectural inductive biases enhances generalization, interpretability, and efficiency, providing a principled framework for AI-driven prediction of complex biological interactions grounded in biophysical mechanisms.

## Results

### 1. Overview of PLMDA-PPI

#### 1.1 Mechanistic inductive biases underlying protein–protein interactions

From a thermodynamic perspective, proteins can transiently encounter each other in the absence of spatiotemporal constraints. However, functional PPIs are distinguished by their ability to adopt specific, low-free-energy configurations corresponding to the native state, forming well-defined basins on the free-energy landscape. In contrast, non-functional protein collisions fail to settle into stable configurations and thus produce a landscape lacking prominent energetic minima. If both types of protein pairs were simulated over long timescales and their interfacial residue contact propensities estimated, functional PPIs would exhibit distinct, high-probability contact signatures, whereas non-interacting pairs would lack discernible contact signals (Figure 1a). These residue-level interaction patterns—drawn from both interacting and non-interacting protein pairs—encode the mechanistic principles that govern protein associations and furnish a natural, mechanistically grounded inductive bias that guides predictive models toward more robust generalization.

**Figure 1.**
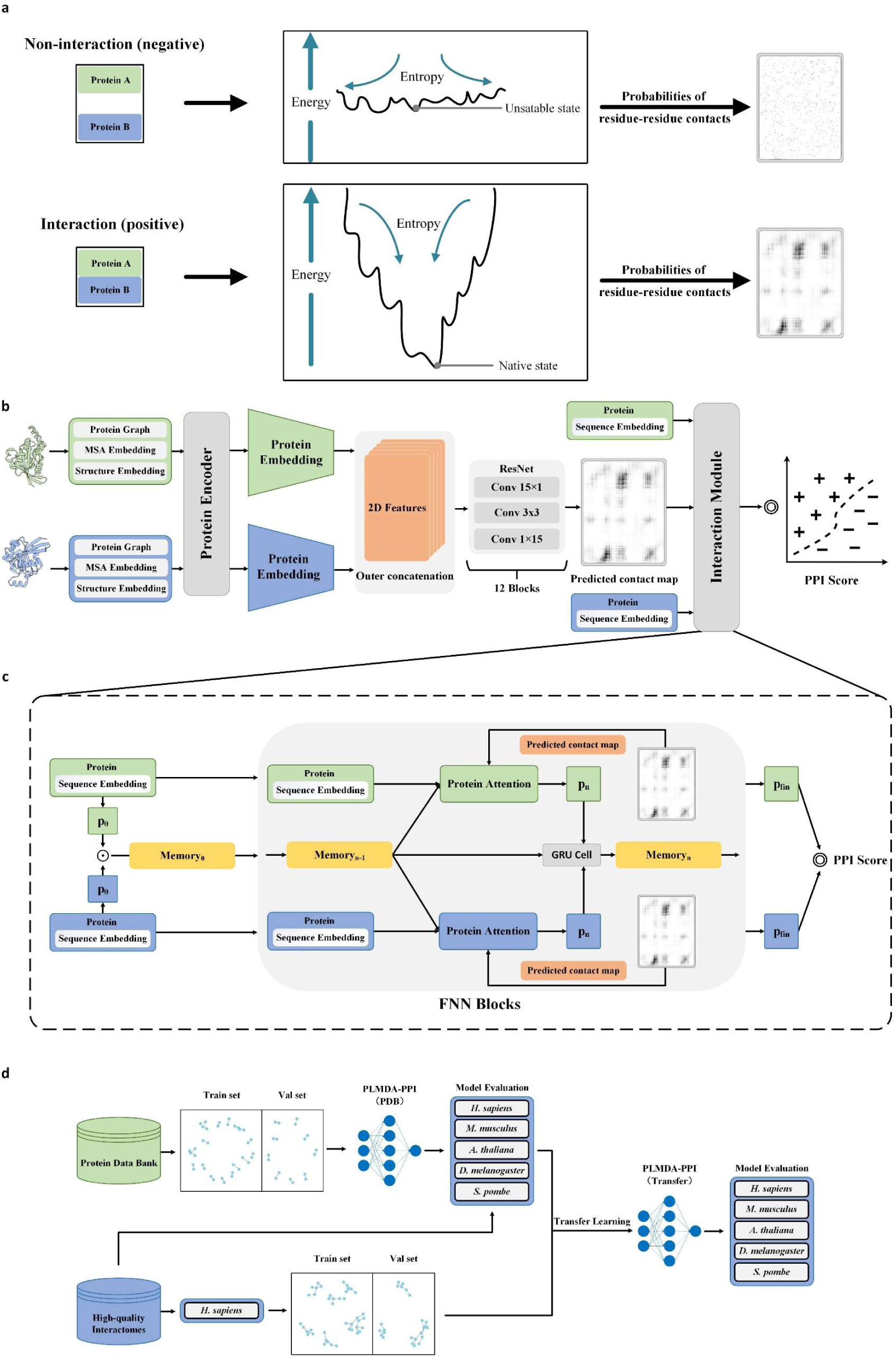
The theoretical basis, model architecture, and training protocol of PLMDA-PPI. (a) Schematic illustration of the mechanistic inductive biases underlying functional protein-protein associations (i.e. interaction) and non-functional collisions (i.e. non-interaction). (b) The overall model architecture of PLMDA-PPI. (c) The interaction prediction module of PLMDA-PPI. (d) The workflow for training and evaluation of PLMDA-PPI.

#### 1.2 Model architecture of PLMDA-PPI incorporating mechanistic inductive biases

Building on the mechanistic inductive biases identified in residue-level contact patterns, we developed PLMDA-PPI, a model explicitly designed to embed these priors for accurate PPI prediction (Figures 1b-c). PLMDA-PPI takes protein sequences, structural information, and multiple sequence alignments (MSAs) as input. From the structures and MSAs, protein language model–embedded geometric graphs are constructed to encode evolutionary and structural constraints. A representation for each protein pair is then created by performing an outer concatenation of their embeddings, which is processed through a dimensional hybrid residual network^33, 34^ to generate a predicted inter-protein contact map. The resulting contact map captures residue-level interaction patterns, producing contact maps with clear, stable residue contacts for functional PPIs and no detectable signals for non-interacting protein pairs. Finally, this contact map guides a dual-attention module^17, 18^ that integrates sequence embeddings from ESM-1b^35^ to generate the final PPI score (Figure 1c). Detailed methods are provided in the Methods section.

#### 1.3 Training and evaluation protocols of PLMDA-PPI

In Figure 1d, we show the protocols we used to train and evaluate PLMDA-PPI. Specifically, PLMDA-PPI was first trained on a non-redundant, structure-informed PPI dataset from the PDB, with the inter-protein contact prediction and PPI prediction modules trained simultaneously using a linear combination of contact loss and PPI loss, allowing the model to leverage residue-level mechanistic signals. The dataset was divided into training and validation sets using a 40% sequence identity threshold at the monomer level, so that no individual protein in the training set exceeds 40% sequence identity with any protein in the validation set, minimizing overfitting. Due to the limited number of non-redundant PDB PPIs, no separate test set was created. To evaluate generalization, the trained model was tested directly on literature-curated, high-quality binary PPIs from HINT for *Homo sapiens* (*H. sapiens*)*, Mus musculus* (*M. musculus*)*, Arabidopsis thaliana* (*A. thaliana*)*, Drosophila melanogaster* (*D. melanogaster*), and *Schizosaccharomyces pombe* (*S. pombe*). Considering differences in collection criteria between the PDB and HINT, the model was further fine-tuned on *H. sapiens* PPIs from HINT via transfer learning. Specifically, in the fine-tuning, the *H. sapiens* dataset was further divided into training, validation, and test sets using a 40% sequence identity cutoff at the monomer level. The *H. sapiens* test set, together with the HINT datasets from other species, was used to assess performance.

For PDB PPIs, input features were derived from experimentally determined monomer structures, whereas for HINT PPIs, where experimental structures are not always available, predicted monomer models from AlphaFoldDB^36^ were used. Negative samples (non-interacting protein pairs) were generated by randomly pairing monomers within each dataset, excluding pairs sharing interacting partners, with a 10:1 negative-to-positive ratio. A detailed description of the training and evaluation procedures is provided in the Methods section.

### 2. PLMDA-PPI performance following initial training on PDB PPIs

#### 2.1 Validation on PDB PPIs

Following initial training on structure-informed PPIs from the PDB, we evaluated PLMDA-PPI on the corresponding validation set for two tasks: (1) PPI prediction as the primary objective, and (2) inter-protein contact prediction (for positive samples only) as an auxiliary task.

For comparison, four reference models—D-SCRIPT, Topsy-Turvy, TT3D, and TUnA—were trained on the same dataset. D-SCRIPT^22^, Topsy-Turvy^23^, and TT3D^24^ include intermediate modules for contact map prediction but differ in input features and training strategies: D-SCRIPT is purely sequence-based, Topsy-Turvy incorporates network-derived indices, and TT3D further integrates a 21-letter structural alphabet (3Di) via Foldseek^37^. These models compute PPI scores using max-pooling over predicted contact maps and rely on sparsity-inducing regularization without explicit supervision from ground-truth contacts. In contrast, PLMDA-PPI explicitly incorporates mechanistic inductive biases by (i) employing a multi-task loss to directly supervise the contact prediction module using experimentally derived inter-protein contacts, and (ii) fusing the predicted residue-level contact maps with protein language model embeddings through a dual-attention network.

Among the reference models, TUnA is a recently developed Transformer-based method that integrates the Spectral-normalized Neural Gaussian Process (SNGP) for uncertainty estimation, enabling it to identify out-of-distribution samples and mitigate overfitting^25^. Unlike PLMDA-PPI, D-SCRIPT, Topsy-Turvy, and TT3D, TUnA does not include an intermediate module for inter-protein contact map prediction, highlighting a key distinction in the utilization of mechanistic information.

In Table 1, we present the performance of PLMDA-PPI and the reference models on the validation set of structure-informed PPIs, considering both PPI prediction and inter-protein contact prediction. For PPI prediction, we report the Area Under the Precision-Recall Curve (AUPRC) and the Area Under the Receiver Operating Characteristic Curve (AUROC). For contact prediction, we evaluate the mean precision of the top 50 predicted residue-residue contacts. Given the 10:1 negative-to-positive sample ratio, a null model would achieve an AUPRC of 0.1 and an AUROC of 0.5. Since AUPRC is more informative for imbalanced datasets^38^, the Precision-Recall curves for all models are additionally visualized in Figure 2a.

**Figure 2.**
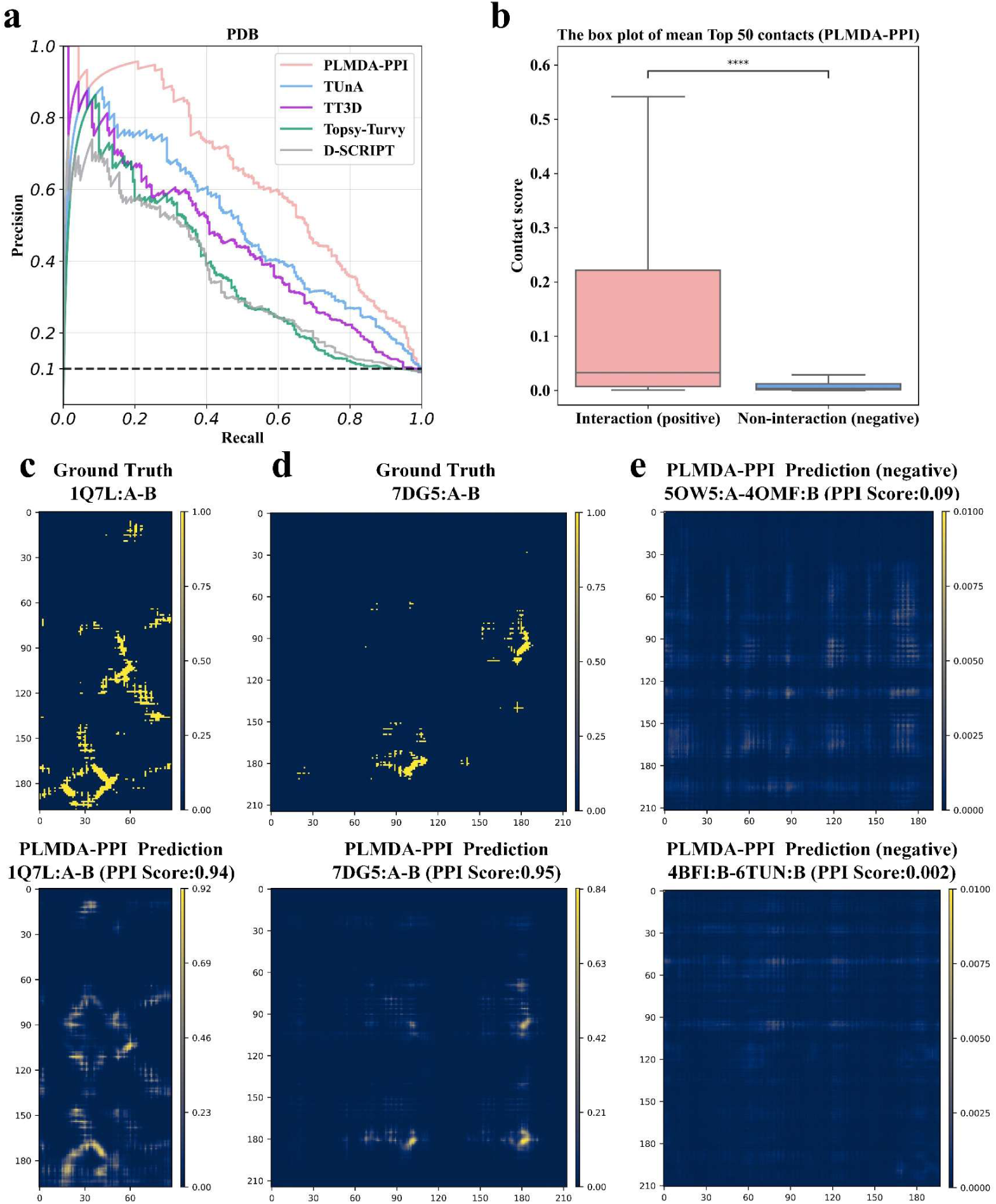
Model validation after initial training on the structure-informed PPI dataset from PDB. (a) The Precision-Recall curves of PLMDA-PPI, D-SCRIPT, Topsy-Turvy, TT3D, and TUnA on the validation set of structure-informed PPIs. The dashed line denotes the expected performance of a null model, with an AUPRC of 0.1. (b) Comparison of the distribution of the mean contact score of the top 50 residue pairs predicted by PLMDA-PPI for interacting and non-interacting protein pairs. Asterisks indicate statistically significant difference based on the Mann-Whitney U test *p*-value: \*\*\*\**p*≤0.0001 (*p*=2.8e-43). (c)∼(d) The experimental (upper panels) and the PLMDA-PPI predicted (lower panels) inter-protein contact maps for two positive samples ((c) PDB 1Q7L:A-B; (d) PDB 7DG5:A-B). (e) The PLMDA-PPI predicted inter-protein contact maps for two negative samples (upper panel: PDB 5OW5:A-4QMF:B; lower panel: PDB 4BFI:B-6TUN:B)

**Table 1.** Validation of the models trained from structure-informed PPIs from PDB.

| Dataset | Model | AUPRC | AUROC | Mean Precision<br>(Top 50 Contact Prediction) |
| --- | --- | --- | --- | --- |
| PDB dataset | D-SCRIPT | 0.324 | 0.684 | 0.026 |
|  | Topsy-Turvy | 0.352 | 0.756 | 0.029 |
|  | TT3D | 0.448 | 0.826 | 0.036 |
|  | TUnA | 0.496 | 0.873 | - |
|  | PLMDA-PPI | **0.637** | **0.911** | **0.287** |

We show performances of PLMDA-PPI, D-SCRIPT, Topsy-Turvy, TT3D, and TUnA on the validation set of structure-informed PPIs from PDB after initial learning. AUPRC: area under precision-recall curve; AUROC: area under receiver operating characteristic curve; Mean Precision (Top 50 Contact Prediction): the mean precisions of the top 50 predicted contacts. TUnA does not predict inter-protein contacts. Top-performing metrics are highlighted in bold.

As shown in Table 1 and Figure 2a, PLMDA-PPI achieves an AUPRC of 0.637 and AUROC of 0.911 for PPI prediction, substantially outperforming all reference models. For inter-protein contact prediction, it reaches a mean precision of 0.287 for the top 50 predicted residue pairs, clearly surpassing the other models, which exhibit minimal contact prediction capability (TUnA does not predict contacts). It is worth noting that inter-protein contact prediction functions only as an auxiliary objective within an intermediate module, primarily introduced to strengthen the generalization of PPI prediction. Therefore, it is understandable that its performance in inter-protein contact prediction does not match that of specialized models (e.g., our previous work PLMGraph-Inter^19^) specifically designed for this task. Figure 2b shows that interacting protein pairs exhibit significantly stronger predicted contact signals than non-interacting pairs, validating that residue-level interactions are effectively leveraged. Examples of experimental and predicted contact maps are shown in Figures 2c–e. Specifically, positive samples correctly predicted as interacting exhibit clear residue–residue contact signals largely consistent with experimentally determined contacts (Figures 2c-d). In contrast, negative samples correctly predicted as non-interacting show much weaker contact signals (Figures 2e).

#### 2.2 Evaluation on multi-species HINT PPIs

We further evaluated PLMDA-PPI and the four reference models on high-quality, literature-curated binary PPIs from the HINT database for *H. sapiens*, *M. musculus*, *A. thaliana*, *D. melanogaster*, and *S. pombe*. Since most HINT PPIs lack structural information, evaluation focused solely on PPI prediction. As shown in Table 2 and Figures 3a–e, PLMDA-PPI consistently outperformed the reference models across all species (also see Figure S1). In addition, we examined residue-level contact probability maps across these datasets, finding that positive samples display significantly stronger residue contact signals than negative ones (Figure S2), consistent with our observations on the PDB PPI dataset and indicating that residue contact signals effectively guide PPI predictions on the HINT datasets as well.

**Figure 3.**
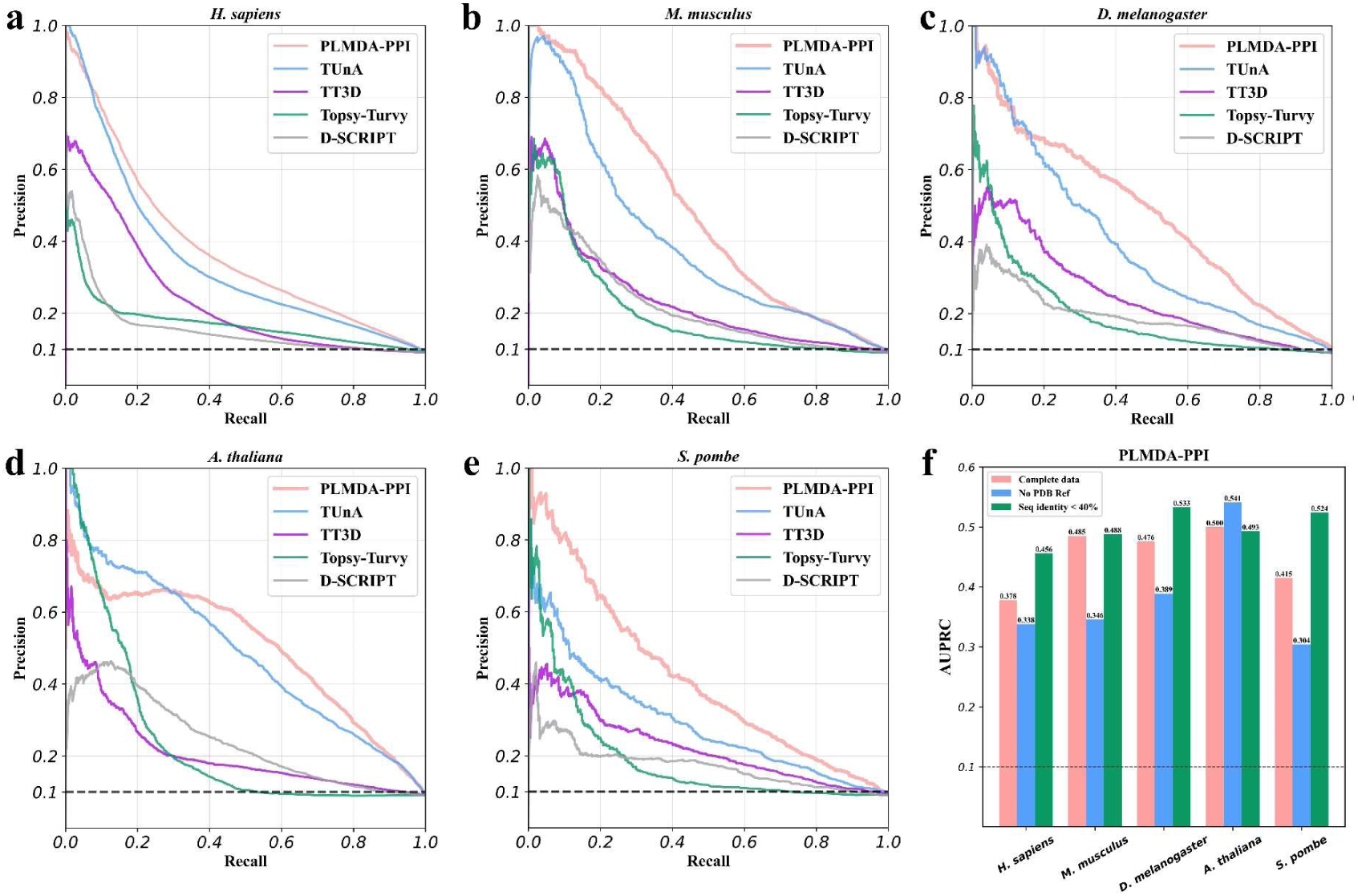
Performances of PLMDA-PPI, D-SCRIPT, Topsy-Turvy, TT3D, and TUnA on the HINT datasets of multiple species after initial training on the structure-informed PPI dataset from PDB. (a)∼(e) The Precision-Recall curves on *H. sapiens* (a), *M. musculus* (b), *D. melanogaster* (c), *A. thaliana* (d), and *S. pombe* (e). (f) Performance variations of PLMDA-PPI (in terms of AUPRC) on the HINT datasets after using different criteria to filter the datasets. The dashed lines denote the expected performance of a null model, with an AUPRC of 0.1.

**Table 2.**
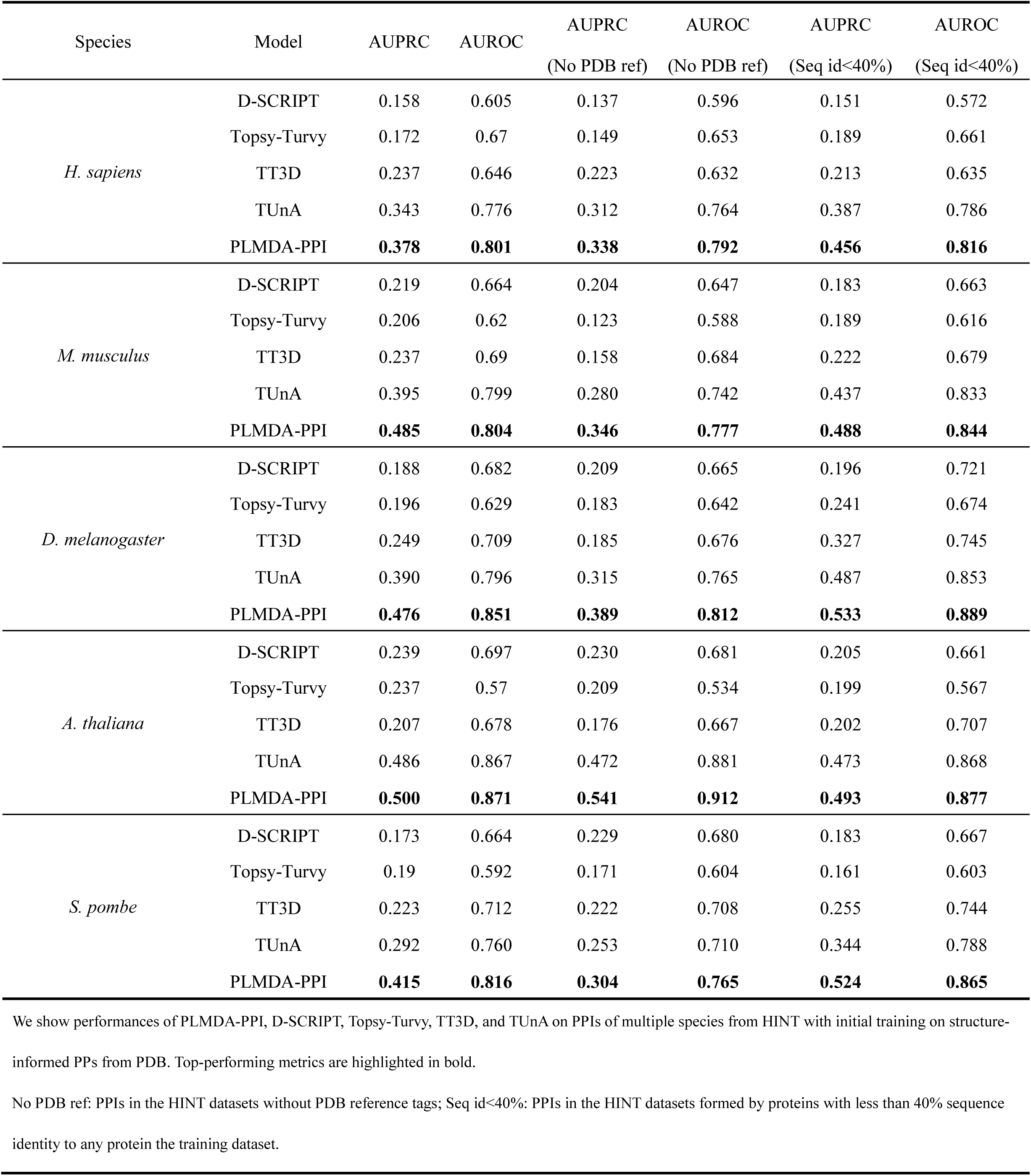
Evaluation of the models trained from PDB PPIs on multiple species PPIs from HINT database.

To assess the model’s ability to predict interactions without structural references, we excluded all HINT interactions with any PDB entry and generated new negative samples at a 10:1 ratio per species. We found that the predictive performance of PLMDA-PPI declined moderately across datasets other than *A. thaliana*, and the other four reference models exhibited similar trends, indicating that models trained solely on PDB-based data may not generalize well to structurally unresolved PPIs (Table 2, Figure 3f). The results highlight the importance of incorporating structure-independent PPI data to further improve predictive performance.

To further evaluate generalization to novel proteins, we applied an additional filtering step, excluding all PPIs involving proteins sharing >40% sequence identity with training set proteins and regenerated negative samples at a 10:1 ratio. PLMDA-PPI showed no systematic performance degradation; modest improvements were even observed for *H. sapiens*, *D. melanogaster*, and *S. pombe*. Similar trends were also observed for the reference models (Table 2, Figure 3f). This is consistent with the fact that the PDB and HINT datasets differ substantially in their protein coverage. The performance gains may result from filtering out particularly challenging prediction cases. Overall, these results demonstrate that PLMDA-PPI generalizes effectively across species and to previously unseen proteins, regardless of sequence similarity.

### 3. Performance of PLMDA-PPI after fine-tuning on HINT *H. sapiens* PPIs

#### 3.1 Evaluation on multi-species HINT PPIs

Given the discrepancy in data collection criteria between PDB and HINT, we adopted a transfer learning strategy to enhance the performance of our models in predicting PPIs under the HINT standard. Specifically, we fine-tuned the models initially trained on PDB data—including PLMDA-PPI and four reference models—using the *H. sapiens* subset from the HINT datasets. To evaluate their performance after fine-tuning, we tested the models on the *H. sapiens* test set, as well as on datasets from *M. musculus*, *A. thaliana, D. melanogaster*, and *S. pombe* in HINT. As shown in Table 3 and Figures 4a–e, PLMDA-PPI consistently outperformed the four reference models across all five datasets (also see Figure S3), and transfer learning led to a substantial improvement in its predictive performance compared to the pre-transfer version (Figure 4f). Moreover, analysis of residue-level contact predictions showed that positive samples continue to exhibit significantly stronger signals than negative ones (Figure S4), indicating that residue-level contact signals remain instrumental in guiding PPI predictions.

**Figure 4.**
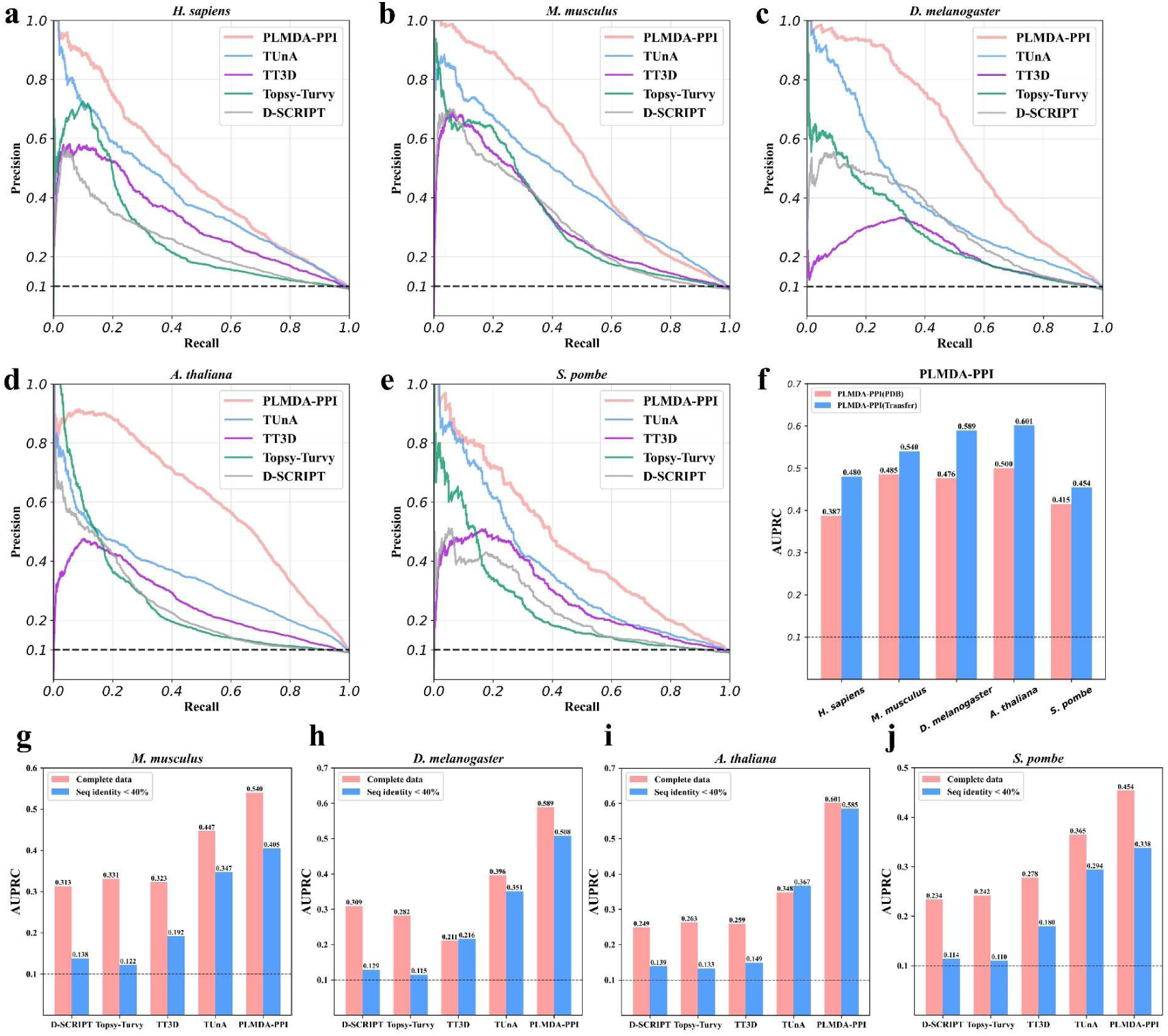
Performances of PLMDA-PPI, D-SCRIPT, Topsy-Turvy, TT3D, and TUnA on the HINT PPI datasets of multiple species after transfer learning on the HINT *H. sapiens* PPI dataset. (a)∼(e) The Precision-Recall curves on *H. sapiens* (test set) (a), *M. musculus* (b), *D. melanogaster* (c), *A. thaliana* (d), and *S. pombe* (e). (f) Performance comparison of PLMDA-PPI (in terms of AUPRC) on the HINT datasets before and after transfer learning. (g)∼(j) Performance variations (in terms of AUPRC) of PLMDA-PPI, D-SCRIPT, Topsy-Turvy, TT3D, and TUnA on the HINT PPI datasets of *M. musculus* (g), *D. melanogaster* (h), *A. thaliana* (i), and *S. pombe* (j) after excluding all PPIs involving proteins with >40% sequence identity to any protein in the *H. sapiens* training set. The dashed line denotes the expected performance of a null model, with an AUPRC of 0.1.

**Table 3.** Evaluation of the HINT fine-tuned models on multiple species PPIs from HINT database.

| Species | Model | AUPRC | AUROC | AUPRC<br>(Seq id < 40%) | AUROC<br>(Seq id < 40%) |
| --- | --- | --- | --- | --- | --- |
| *H. sapiens*<br>(test set) | D-SCRIPT | - | - | 0.245 | 0.713 |
|  | Topsy-Turvy | - | - | 0.273 | 0.696 |
|  | TT3D | - | - | 0.318 | 0.784 |
|  | TUnA | - | - | 0.409 | 0.792 |
|  | PLMDA-PPI | - | - | **0.480** | **0.840** |
| *M. musculus* | D-SCRIPT | 0.313 | 0.719 | 0.138 | 0.661 |
|  | Topsy-Turvy | 0.331 | 0.732 | 0.122 | 0.583 |
|  | TT3D | 0.323 | 0.755 | 0.192 | 0.656 |
|  | TUnA | 0.447 | 0.844 | 0.347 | 0.795 |
|  | PLMDA-PPI | **0.540** | **0.840** | **0.405** | **0.801** |
| *D. melanogaster* | D-SCRIPT | 0.309 | 0.67 | 0.129 | 0.643 |
|  | Topsy-Turvy | 0.282 | 0.671 | 0.115 | 0.559 |
|  | TT3D | 0.211 | 0.743 | 0.216 | 0.665 |
|  | TUnA | 0.396 | 0.808 | 0.351 | 0.809 |
|  | PLMDA-PPI | **0.589** | **0.871** | **0.508** | **0.855** |
| *A. thaliana* | D-SCRIPT | 0.249 | 0.754 | 0.139 | 0.637 |
|  | Topsy-Turvy | 0.263 | 0.721 | 0.133 | 0.578 |
|  | TT3D | 0.259 | 0.724 | 0.149 | 0.593 |
|  | TUnA | 0.348 | 0.818 | 0.367 | 0.824 |
|  | PLMDA-PPI | **0.601** | **0.898** | **0.585** | **0.899** |
| *S. pombe* | D-SCRIPT | 0.234 | 0.682 | 0.114 | 0.570 |
|  | Topsy-Turvy | 0.242 | 0.664 | 0.110 | 0.537 |
|  | TT3D | 0.278 | 0.744 | 0.180 | 0.625 |
|  | TUnA | 0.365 | 0.770 | 0.294 | 0.759 |
|  | PLMDA-PPI | **0.454** | **0.830** | **0.338** | **0.826** |

We show performances of PLMDA-PPI, D-SCRIPT, Topsy-Turvy, TT3D, and TUnA on PPIs of multiple species from HINT after transfer learning on the HINT *H. sapiens* PPI dataset. Top-performing metrics are highlighted in bold.
Seq id<40%: PPIs in the HINT datasets formed by proteins with less than 40% sequence identity to any protein the training dataset.

#### 3.2 Evaluation on proteins with low sequence similarity to training data

In the previous evaluations, only the *H. sapiens* test set was strictly filtered to exclude any PPIs involving proteins with more than 40% sequence identity to those in the transfer learning training set. For PPIs from other species, no such sequence-based filtering was applied, leaving open the possibility that models could partially rely on sequence similarity when making predictions. To provide a more rigorous assessment of model generalization to novel proteins, we applied stringent filtering across all HINT datasets. Specifically, we removed any PPIs containing proteins with greater than 40% sequence identity to proteins in the *H. sapiens* training set. Negative samples were re-generated for each species at a 10:1 ratio relative to positive samples. These filtered datasets were then used to re-evaluate model performance, as shown in Table 3 and Figures 4g–j.

Under this more stringent evaluation, all models—including PLMDA-PPI and the four reference models—experienced some performance decline on the filtered datasets. This drop may reflect evolutionary relationships between PPIs of non-*H. sapiens* species and *H. sapiens,* which can allow models to leverage sequence similarity. Nevertheless, PLMDA-PPI retained substantial predictive power, whereas D-SCRIPT, Topsy-Turvy, and TT3D showed sharp declines, approaching null-model performance (AUPRC = 0.1), consistent with the previous observation^14^. TUnA exhibited a moderate decline, yet it still outperformed these three reference models while remaining noticeably inferior to PLMDA-PPI. Overall, these results further underscore that PLMDA-PPI demonstrates markedly superior generalization to previously unseen proteins compared to existing methods.

We also assessed the pre-transfer learning model on the same filtered datasets. Across most species, the transfer-learned model substantially outperformed the pre-transfer version. The only exception was the *S. pombe* dataset, where AUPRC slightly decreased (0.342 → 0.338), though AUROC improved (0.794 → 0.826; Supplementary Table S1). These findings indicate that the observed gains from transfer learning cannot be attributed solely to high sequence similarity between training and test proteins, supporting the broader generalization benefit of the fine-tuning procedure.

#### 3.3 Evaluation of HINT fine-tuned models on PDB PPIs

To evaluate the generalization of transfer-learned models, we tested whether models fine-tuned on the HINT PPIs retained predictive performance on the original PDB-based structure-informed PPIs. We applied the transfer-learned models to the previously used validation set of PDB PPIs. Table 4 summarizes the results for both the primary task of PPI prediction and the auxiliary task of inter-protein contact prediction. PLMDA-PPI not only outperforms all reference models in both tasks but, when compared with the initial model, the fine-tuning even slightly improves its PPI prediction performance (Figure 5a). In contrast, the other models experience substantial accuracy declines (Figures 5b–e). For contact prediction, PLMDA-PPI exhibits a moderate decrease in precision for the top 50 predicted contacts (0.287 → 0.230), but still maintains significant predictive power (also see Figure S5), whereas reference models show virtually no capacity to predict residue-level contacts. Moreover, analysis of the inter-protein contact predictions from PLMDA-PPI showed that the interacting protein pairs still exhibit significantly stronger inter-protein signals compared with non-interacting pairs (Figure 5f). These results underscore the superior generalization capability of PLMDA-PPI after transfer learning.

**Figure 5.**
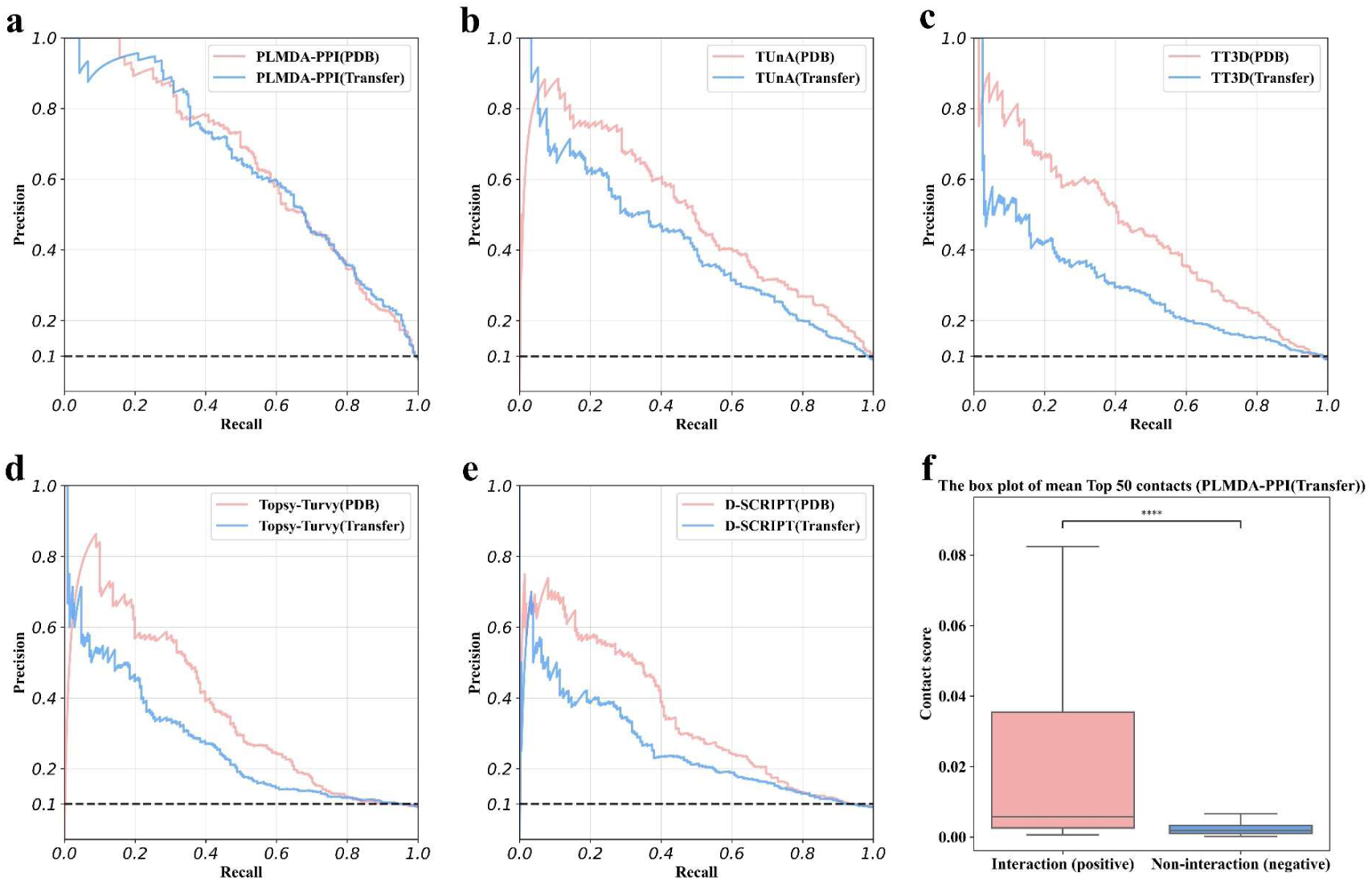
Re-evaluation of the models after transfer learning–based fine-tuning using the HINT *H. sapiens* PPI dataset on the validation set of structure-informed PPIs derived from PDB. Comparisons of the performances (using Precision-Recall curves) before and after transfer learning for PLMDA-PPI (a), TUnA (b), TT3D (c), Topsy-Turvy (d), and D-SCRIPT (e). (f) Comparison of the distribution of the mean contact score of the top 50 residue pairs predicted by PLMDA-PPI after transfer learning for interacting and non-interacting protein pairs. Asterisks indicate statistically significant difference based on the Mann-Whitney U test *p*-value: \*\*\*\**p*≤0.0001 (*p*=1.1e-42).

**Table 4.** Evaluation of the HINT fine-tuned models on PDB PPIs.

| Dataset | Model | AUPRC | AUROC | Mean Precision<br>(Top 50 Contact Prediction) |
| --- | --- | --- | --- | --- |
| PDB dataset | D-SCRIPT | 0.258 | 0.719 | 0.026 |
|  | Topsy-Turvy | 0.271 | 0.697 | 0.029 |
|  | TT3D | 0.296 | 0.757 | 0.036 |
|  | TUnA | 0.421 | 0.827 | - |
|  | PLMDA-PPI | **0.642** | **0.907** | **0.230** |

We re-evaluated the performance of PLMDA-PPI, D-SCRIPT, Topsy-Turvy, TT3D, and TUnA after transfer learning on the HINT *H. sapiens* PPI dataset, using the validation set of structure-informed PPIs derived from PDB. Top-performing metrics are highlighted in bold.

### 4. Benchmarking PLMDA-PPI against AF2Complex, RF2-Lite, and RF2-PPI

We next conducted a systematic benchmarking of PLMDA-PPI against several recently developed PPI prediction models built upon the landmark protein structure prediction frameworks AlphaFold2 and RoseTTAFold2, including AF2Complex, RF2-Lite, and RF2-PPI. AF2Complex, based on AlphaFold-Multimer^31^, introduces a specialized interface score (iScore) to evaluate potential PPIs. However, the extremely high computational cost of AlphaFold-Multimer renders AF2Complex impractical for large-scale PPI screening. RF2-Lite and RF2-PPI are streamlined versions of RoseTTAFold2 optimized for inference speed and calibrated specifically for bacterial and human PPIs, respectively. Nonetheless, both models still incur substantially higher computational costs than PLMDA-PPI and require generating paired MSAs containing inter-protein co-evolutionary information for each candidate pair, further increasing the computational burden. In contrast, PLMDA-PPI only requires one-time feature extraction for individual protein monomers, greatly reducing resource demands.

To benchmark these methods across five representative species, we randomly selected 50 positive PPIs per species and obtained tenfold negative samples (500 non-interacting protein pairs) by randomly drawing from the existing negative dataset, due to the substantial inference time of these methods. Notably, RF2-Lite and RF2-PPI originally relied on species-specific proteome/genome databases to produce paired MSAs, which are not generalizable; we therefore employed ColabFold^39^ to generate paired MSAs. Besides, unlike the lightweight models, retraining any of these models on the same training set as PLMDA-PPI was practically infeasible given the much large datasets and computational resources required, so all three models were evaluated using their publicly available parameters.

Evaluation results on HINT-derived subsets are shown in Figures 6a–e and Table 5. PLMDA-PPI achieves the highest predictive performance in four out of five species; the only exception is *S. pombe*, where it outperforms RF2-Lite but is outperformed by AF2Complex and RF2-PPI (also see Figure S6). For AF2Complex, two AlphaFold-Multimer weight sets were tested: model_1_multimer_v3 (with monomer templates) and model_3_multimer_v3 (without monomer templates), denoted AF2Complex(1) and AF2Complex(3). Inference times per PPI excluding the time for input feature generation on a single NVIDIA A6000 GPU are summarized in Figure 6f. PLMDA-PPI averages ∼0.24 s per prediction—30–50 times faster than RF2-Lite (12.8 s) and RF2-PPI (7.8 s), and 1,000–2,000 times faster than AF2Complex(1) (535.4 s) and AF2Complex(3) (285.8 s). Furthermore, unlike the other methods, PLMDA-PPI only needs one-time feature preparation for individual protein monomers and does not require generating paired MSAs for each protein pair, further amplifying its computational advantage in large-scale PPI screening.

**Figure 6.**
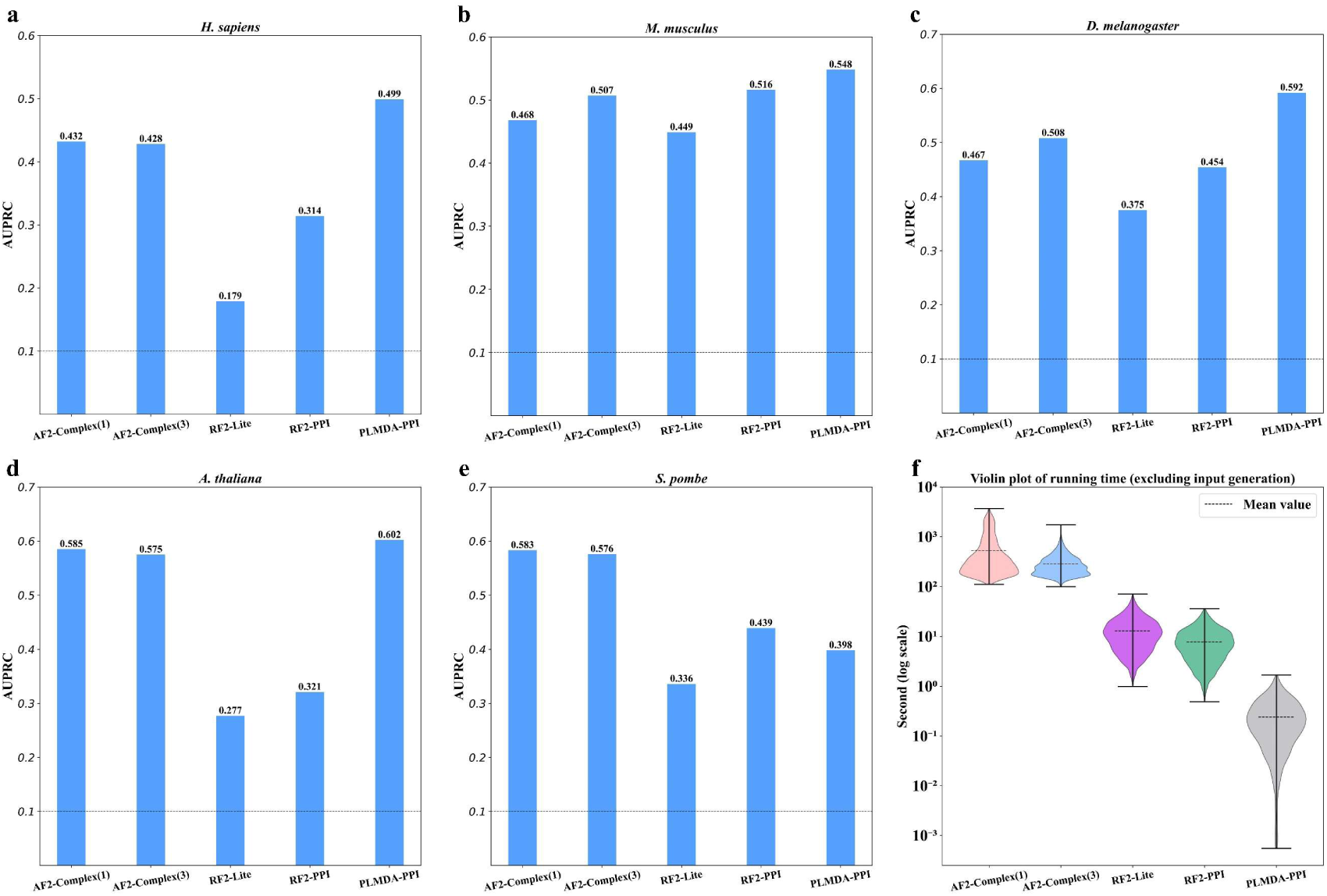
Performances of PLMDA-PPI, AF2-Complex, RF2-Lite, and RF2-PPI on subsets of PPIs sampled from HINT datasets for *H. sapiens* (a), *M. musculus* (b), *D. melanogaster* (c), *A. thaliana* (d), and *S. pombe* (e). For each species, 50 PPIs were randomly selected as positive samples, and ten times as many (i.e., 500) non-interacting protein pairs were generated through random pairing as negative samples. (f) Violin plots of inference-only running times for AF2-Complex, RF2-PPI, and RF2-Lite (time for input feature generation excluded). For AF2Complex, two sets of AlphaFold-Multimer model weight: model_1_multimer_v3 (using monomer templates) and model_3_multimer_v3 (without monomer templates) were employed—denoted as AF2Complex(1) and AF2Complex(3), respectively. The dashed line denotes the expected performance of a null model, with an AUPRC of 0.1.

**Table 5.** Performances of AF2-Complex, RF2-Lite, RF2-PPI, and PLMDA-PPI on multiple species PPIs from HINT database.

| Species | Model | AUPRC | AUROC |
| --- | --- | --- | --- |
| *H. sapiens* | AF2-Complex(1) | 0.432 | 0.696 |
|  | AF2-Complex(3) | 0.428 | 0.702 |
|  | RF2-Lite | 0.179 | 0.594 |
|  | RF2-PPI | 0.314 | 0.645 |
|  | PLMDA-PPI | **0.499** | **0.839** |
| *M. musculus* | AF2-Complex(1) | 0.468 | 0.687 |
|  | AF2-Complex(3) | 0.507 | 0.734 |
|  | RF2-Lite | 0.449 | 0.717 |
|  | RF2-PPI | 0.516 | 0.727 |
|  | PLMDA-PPI | **0.548** | **0.877** |
| *D. melanogaster* | AF2-Complex(1) | 0.467 | 0.748 |
|  | AF2-Complex(3) | 0.508 | 0.753 |
|  | RF2-Lite | 0.375 | 0.716 |
|  | RF2-PPI | 0.454 | 0.675 |
|  | PLMDA-PPI | **0.592** | **0.861** |
| *A. thaliana* | AF2-Complex(1) | 0.585 | 0.729 |
|  | AF2-Complex(3) | 0.575 | 0.766 |
|  | RF2-Lite | 0.277 | 0.689 |
|  | RF2-PPI | 0.321 | 0.631 |
|  | PLMDA-PPI | **0.602** | **0.906** |
| *S. pombe* | AF2-Complex(1) | **0.583** | 0.762 |
|  | AF2-Complex(3) | 0.576 | 0.757 |
|  | RF2-Lite | 0.336 | 0.711 |
|  | RF2-PPI | 0.439 | 0.665 |
|  | PLMDA-PPI | 0.398 | **0.818** |

We show performances of AF2Complex, RF2-Lite, RF2-PPI, and PLMDA-PPI on the subsets of PPIs sampled from HINT datasets. For AF2Complex, we employed two sets of AlphaFold-Multimer model weights recommended in the documentation: model\_1\_multimer\_v3 (using monomer templates) and model\_3\_multimer\_v3 (without monomer templates)—denoted as AF2Complex(1) and AF2Complex(3), respectively. Top-performing metrics are highlighted in bold.

Collectively, these results demonstrate that PLMDA-PPI combines high predictive performance with exceptional computational efficiency, highlighting that inter-protein co-evolutionary information from paired MSAs is not indispensable for accurate PPI prediction.

### 5. Ablation studies

We further conducted ablation studies to systematically assess the contributions of the key components and training strategies of PLMDA-PPI, covering both the architectural design and training approach. Specifically, we first retrained the model after removing major modules using the structure-informed PPI training set derived from PDB and evaluated performance on the corresponding validation set. When we removed the MSA and structural protein language model representations embedded in the geometric graphs—retaining only the geometric graph—and replaced the sequence-based protein language model embeddings in the interaction prediction module with geometric node features, the model’s performance dropped substantially (AUPRC decreased from 0.637 to 0.229). Restoring the protein representations in the interaction module to language model embeddings led to a significant performance recovery (AUPRC = 0.593), although still inferior to the full model. These results highlight the crucial role of language model representations in our framework. Additionally, replacing the dual-attention network in our model with the customized max pooling module used in D-SCRIPT, Topsy-Turvy, and TT3D also led to a notable performance decline (AUPRC = 0.411), underscoring the importance of the dual-attention network module (Figure 7a).

**Figure 7.**
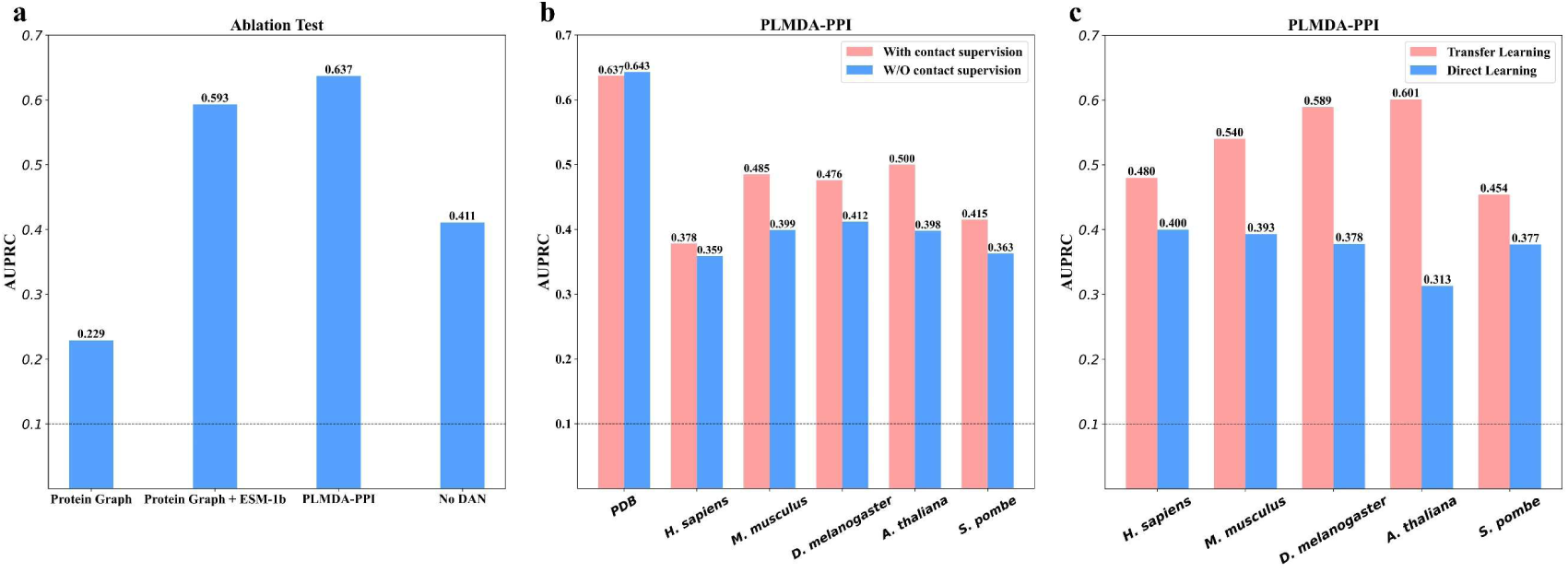
Ablation studies for PLMDA-PPI. (a) Performance variations (in terms of AUPRC) on the validation set of structure-informed PPIs when key components are removed from the model architecture. (b) Performance comparison (in terms of AUPRC) between models trained with and without inter-protein contact supervision. (c) Performance comparison (in terms of AUPRC) between models with and without the incorporation of structure-informed PPIs from the PDB for the initial training. The dashed line denotes the expected performance of a null model, with an AUPRC of 0.1.

To further probe the impact of inductive bias from inter-protein contact supervision, we trained the model without the multitask loss for residue-residue contact prediction, optimizing only for the primary interaction prediction task. On the PDB validation set, this change led to negligible improvement (AUPRC = 0.643). However, across the HINT multi-species datasets, performance consistently declined, demonstrating that learning to predict residue-level contacts enhances the model’s generalization to unseen proteins (Figure 7b).

Finally, we trained a model directly on *H. sapiens* PPIs from HINT without any PDB pretraining. Across all species, this model performed substantially worse than the PDB-pretrained and fine-tuned model, further confirming that pretraining on structure-informed datasets establishes inductive biases that are crucial for robust cross-species generalization (Figure 7c and Supplementary Table S2).

Collectively, these ablation studies demonstrate that the inductive biases embedded in the architecture and training strategy—via protein language model embeddings, dual-attention integration, and residue-level contact supervision—are key drivers of PLMDA-PPI’s strong generalization performance.

### Conclusion

In this study, we present PLMDA-PPI, a mechanism-aware framework for protein–protein interaction prediction that incorporates residue-level biophysical inductive biases, derived from protein language model–embedded geometric representations, into a dual-attention architecture. By jointly learning global interaction likelihoods and inter-protein contact patterns, PLMDA-PPI captures mechanistic signatures that distinguish functional PPIs from non-functional encounters, enabling robust generalization beyond the training distribution.

Across structure-informed PDB PPIs, multi-species HINT datasets, and stringent sequence-dissimilarity settings, PLMDA-PPI consistently outperforms recent lightweight deep learning models and maintains predictive power even when sequence similarity cues are minimized. Transfer learning on *H. sapiens* PPIs further enhances model performance while preserving the mechanistic generalization imparted by structure-informed pretraining. Notably, PLMDA-PPI also achieves accuracy comparable to or even exceeding that of structure-prediction model–based methods—including AF2Complex, RF2-Lite, and RF2-PPI—while requiring orders-of-magnitude fewer computational resources and eliminating dependence on paired MSAs.

Ablation studies confirm that the framework’s key components—protein language model representations, dual-attention integration, and residue-level contact supervision—constitute essential inductive biases that substantially improve cross-species and out-of-distribution performance. Together, these findings underscore that incorporating mechanistic priors directly into model architectures provides a principled route toward interpretable, generalizable, and computationally efficient AI for complex biological interaction prediction.

## Methods

### 1. Dataset preparation

#### 1.1 PPI database selection

We collected structure-resolved interactions from the PDB and experimentally validated non-structural PPI datasets for model development and evaluation. Specifically, the PDB provides high-quality multimer structures that directly capture physical contacts but offers limited species-level coverage, motivating the incorporation of additional curated resources. High-throughput datasets, such as HuRi (Y2H)^7^ and BioPlex (AP–MS)^2^, expand coverage but contain substantial noise.

Computationally compiled repositories, including STRING^13^ and HINT^21^, vary widely due to differences in evidence integration. Notably, STRING’s “Physical Interaction” category does not distinguish direct from indirect co-complex associations^13^. In contrast, HINT offers a high-quality subset of literature-curated binary interactions, supported by at least two independent low-throughput experiments across multiple species, making it one of the most reliable sources for direct PPIs^9, 21^. Accordingly, we used PDB-derived PPIs as structure-informed data and the HINT datasets across multiple species for model development and cross-species evaluation.

#### 1.2 Structure-informed PPI collection from PDB

We constructed a structure-informed PPI dataset by filtering PDB entries deposited before October 24, 2022. Entries were retained if they met the following criteria: X-ray diffraction as the experimental method, resolution ≤4 Å, heteromeric protein composition, and ≥2 protein chains, yielding 27217 PDB entries containing 43,983 biological assemblies. Protein pairs were then extracted from biological assemblies, resulting in 438,554 candidate interactions. To remove redundancy, all 877,108 protein sequences were clustered at 40% identity using CD-HIT^40^, and each pair was assigned a cluster-pair label; pairs sharing the same unordered cluster pair were considered redundant. Inter-protein contact maps were computed using a heavy-atom distance cutoff of 8 Å, and only pairs with >50 contacts were kept. For redundant sets, the pair with the highest contact count was retained. Additional filtering removed proteins with chain lengths outside 30–521 residues, structures with >20% missing residues, and self-interactions (proteins from the same cluster). The final dataset contained 6,189 non-redundant structure-informed PPIs comprising 11,119 unique monomers grouped into 5,952 sequence clusters.

#### 1.3 PPI collection for multiple species from HINT

High-quality, literature-curated binary PPIs for *H. sapiens*, *M. musculus*, *A. thaliana*, *D. melanogaster*, and *S. pombe* were obtained from the HINT database. We removed self-interactions, excluded PPIs involving proteins shorter than 30 residues or longer than 1,024 residues, and discarded interactions involving protein isoforms. The resulting datasets comprised 23,311 PPIs for *H. sapiens*, 2,369 for *M. musculus*, 2,789 for *A. thaliana*, 1,336 for *D. melanogaster*, and 791 for *S. pombe*.

### 2. Model construction of PLMDA-PPI

#### 2.1 Input features

##### 2.1.1 Protein geometric graph

Given a protein’s 3D structure, we constructed its geometric graph using the same protocol as in PLMGraph-Inter^19^. Each residue was represented as a node, and edges were added between residues whose Cα–Cα distance was ≤18 Å. For every node and edge, we computed geometric features consisting of inter-atomic distance scalars and orientation vectors derived from the local coordinate frames, ensuring SE(3)-invariant representations. Additional details of the geometric graph formulation are provided in our previous work^19^.

##### 2.1.2 Embeddings of sequence, MSA, and structure

We employed ESM-1b^35^ to generate sequence embeddings and ESM-IF^41^ to produce structure-based embeddings. Multiple sequence alignments (MSAs) were constructed using Jackhmmer^42^ (default parameters) against the UniRef90^43^ database. For each protein, MSA representations were composed of two components: embeddings generated by ESM-MSA-1b^44^ and position-specific scoring matrices (PSSMs) computed from the alignment statistics.

#### 2.2 Protein encoder

Protein encoding followed the protocol of PLMGraph-Inter. For each protein, node-level scalar features were constructed by concatenating MSA-derived embeddings, structure embeddings, and residue-level geometric scalar descriptors. These scalar features, together with vector geometric features, were processed through three layers of GVP-based message passing^45^, which apply rotationally invariant updates to scalar channels and rotationally equivariant updates to vector channels. The updated scalar and vector features were then combined to yield a 1D per-residue representation. As a key modification, PLMDA-PPI does not incorporate sequence embeddings (ESM-1b) into the protein graph; Instead, sequence representations are used exclusively within the interaction prediction module, as our internal evaluations indicated that incorporating them into both the protein graph and the interaction module substantially increased false-positive rates.

#### 2.3 Inter-protein contact prediction module

The inter-protein contact prediction module takes the 1D residue-level representations generated by the protein encoder and models pairwise interactions between residues of two proteins. Given two representations ***A***∈ℝ^**L1**×**d**^ and ***B***∈ℝ^**L2**×**d**^, where ***L_1_*** and ***L_2_*** denote protein lengths and **d**=448 is the embedding dimension), each representation is broadcast along the partner sequence dimension and concatenate to form a joint 2D representation **C**∈ℝ^**2d**×**L**^_**1**_^×**L**^_**2**_ as follows:

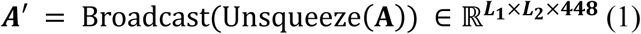

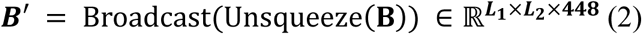

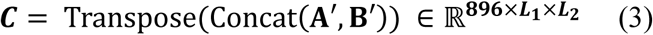

The resulting 2D representation is processed by a dimensional hybrid residual network to predict residue–residue contact probabilities. Specifically, a 1 × 1 convolution first reduces the channel dimension from 896 to 96, followed by 12 successive dimensional hybrid residual blocks^33, 34^. A final 1 × 1 convolution that maps the 96 channels to a single channel, and a sigmoid activation produces the predicted inter-protein contact map.

#### 2.4 Interaction prediction module

PPIs are governed by physicochemical interactions between interfacial residues. Accordingly, interacting protein pairs are expected to display stronger and more specific residue-residue interaction signals in predicted inter-protein contact maps than non-interacting pairs. To leverage this, the contact maps generated by the previous module are used to inform the protein interaction prediction module by integrating single-sequence-based protein language model embeddings (ESM-1b). Only these single-sequence embeddings are used, as including representations existing in the contact map prediction module was found to substantially increase false-positive predictions in our in-house tests. This effect likely arises from the high sequence specificity of PPIs: incorporating explicit structural information or MSA-based embeddings can reduce the model’s sensitivity to sequence-level signals, causing it to erroneously predict interactions for protein pairs with similar monomeric folds to known interacting proteins, even when they do not interact.

We adapted the Dual Attention Network^17, 18^ for multimodal reasoning (r-DAN) to model PPI prediction as a Visual Question Answering (VQA) task. In this framework, protein sequence embeddings replace the visual and textual features in VQA, while the predicted inter-protein contact map serves as a weight matrix guiding information transfer between residues of the two proteins. The architecture of the interaction prediction module is as follows:

First, a linear transformation layer is applied to reduce the dimension of the sequence embeddings generated by ESM-1b from 1280 to 448 (***P***_**1**_ ∈ ℝ*^L^*_1_^×448^, ***P***_**2**_ ∈ ℝ*^L^*_2_^×448^). Then two feed-forward network (FNN) blocks were successively applied to generate the protein context vectors 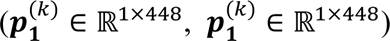 and update the memory vector 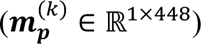 based on the protein embeddings (***P*_1_**, ***P*_2_**) the memory vector 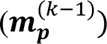 in the previous step, and the predicted inter-protein contact map (***Cmap***) through Equations (4)∼(5):

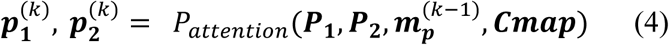

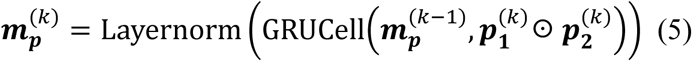

where ☉ represents element-wise multiplication, GRUCell is the computational unit that implements the GRU mechanism for processing sequential data in recurrent neural networks, Layernorm represents Layer Normalization, and *P*_attention_represents Protein Attention, designed to generate protein context vectors by focusing on specific regions in the protein embeddings. Details of this block was described in the following section.

**The Protein Attention block**: First, two intermediate hidden states 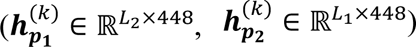 correspond to information transferred from each protein to its partner are calculated using the predicted inter-protein contact map (**Cmap** ∈ ℝ*^L^*_1_^×L2^) as the weight matrix through Equations (6)∼(7):

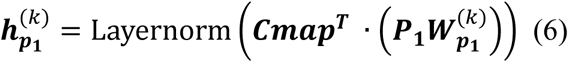

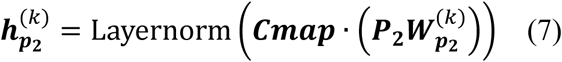

where 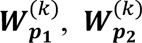 are the network parameters.

Then two protein attention weights 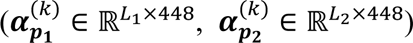 are calculated based on 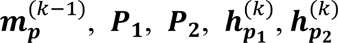 through Equations (8)∼(9):

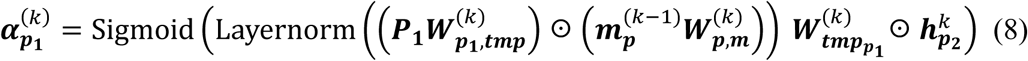

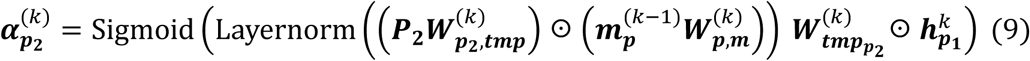

where 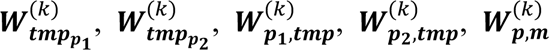 are the network parameters, and Sigmoid represents the Sigmoid Activation.

Finally, the protein context vectors are obtained through first elementwise multiplication of the protein embeddings and the attention weights, followed by summation and average over protein sequences and layer normalization through Equations (10)∼(11):

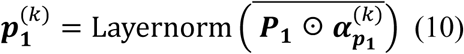

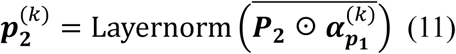

### Memory vector initialization and interaction probability calculation

The initial memory vector 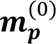 is calculated through Equation (12):

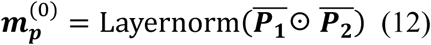

The final predicted interaction probability between two proteins is calculated through Equation (13):

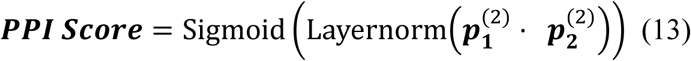

### 3. Training of PLMDA-PPI

#### 3.1 Data leakage in PPI prediction

Previous studies often split PPI datasets randomly into training and test sets. To reduce redundancy, PPIs are typically clustered before splitting: if both proteins in a PPI share sequence similarity above a predefined threshold (commonly 40%) with the corresponding proteins in another PPI, the interactions are considered redundant, and only one representative PPI is retained. This approach partially mitigates data leakage but does not completely eliminate it. Even if protein pairs in the test set do not share sequence similarity with pairs in the training set, individual proteins may still appear in the training data.

Park et al^16^. defined three classes of test sets: C1 (both proteins in test pairs appear in training), C2 (only one protein appears in training), and C3 (no protein overlap). They observed that predictive performance decreases significantly from C1 to C2 and from C2 to C3. Bernett et al^14^. further demonstrated that overlapping proteins between training and test sets can lead models to rely on sequence similarity and node degrees as shortcuts, causing severe overfitting. Minimizing sequence similarity between training and test sets reduces data leakage and reveals that many previously reported high-performing models drop to near-random prediction levels. Therefore, careful handling of data leakage is critical when preparing datasets for model training and evaluation.

#### 3.2 Initial training using structure-informed PPIs from PDB

##### 3.2.1 Dataset preparation

The structure-informed PPI dataset comprises 5,952 sequence clusters derived from protein monomers. These clusters were randomly divided into two non-overlapping groups: Group 1 (5,500 clusters) and Group 2 (452 clusters). Positive training samples consisted of 5,352 PPIs formed between proteins within Group 1, while the validation set included 211 PPIs within Group 2. To prevent potential data leakage, 626 PPIs involving proteins from both groups were excluded. As a result, no proteins in the training or validation sets share sequence similarity above 40%, minimizing overfitting due to redundancy.

Negative samples (non-interacting protein pairs) were generated by randomly pairing monomers within the training and validation sets, after filtering out pairs that shared common interacting partners to avoid including potential true interactions. A 10:1 negative-to-positive ratio was applied for both sets, reflecting the sparsity of true PPIs. Input features were extracted from experimentally determined monomer structures obtained from the corresponding complex structures.

##### 3.2.2 Training protocol

The model was trained using a combined loss function integrating inter-protein contact loss and PPI interaction loss:

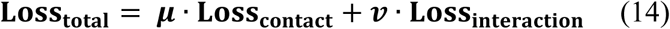

where **μ** = **100** and ***v*** = **1**.

The inter-protein contact loss is defined as:

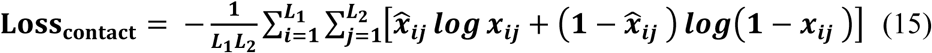

where **L**_**1**_ and **L**_**2**_ are protein lengths, ***x***_***ij***_ represents the contact probability between residue ***i*** and ***j***, and 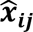 is the ground-truth contact label (set as 1 if contact, and 0 if not). For interacting protein pairs, contact labels were derived from heavy-atom-distances in the experimental structures using an 8Å threshold; for non-interacting proteins pairs, all residue pairs were considered as non-contact 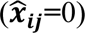.

The interaction loss is defined as:

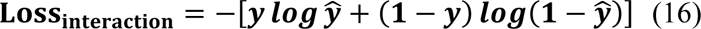

where ***y*** represents the true interaction label (1 for positive pairs, and 0 for negative pairs), and 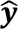 is the predicted interaction probability.

Training was performed using the PyTorch (v2.1.0) on a single NVIDIA A100 GPU. The initial learning rate was set as 1 × 10^-5^, and the AdamW optimizer was used with a batch size of 1. After each epoch, the validation loss was monitored. If the loss did not decrease for 3 consecutive epochs, the learning rate was reduced by a factor of 0.1. After two such reductions, the model weights achieving the lowest validation loss were selected as the final trained model.

#### 3.3 Transfer learning using *H. sapiens* PPIs from HINT database

##### 3.3.1 Datasets for training and validation

To further enhance the model, the *H. sapiens* PPI dataset from the HINT database was used for fine-tuning, initialized with the weights obtained from training on the structure-informed PPI dataset. To minimize the possibility of data leakage in dataset splitting, following the same procedure as previously used in the structure-informed PPI dataset, we first clustered protein monomers in the 23,311 PPIs using CD-HIT with parameters “-c 0.4 -n 0.2 -g 1”, grouping sequences at a 40% identity threshold, which yielded 5873 clusters. The 5873 clusters were then randomly categorized into three groups: group I containing 3800 clusters, group II containing 700 clusters, and group III containing 1373 clusters. We retained only the PPIs formed by proteins within each group and discarded the rest. In addition, to reduce redundancy, for PPIs belonging to the same cluster pair, we randomly selected one representative interaction. As a result, we obtained 4,585 PPIs formed by proteins within group I, 345 PPIs within group II, and 1,502 PPIs within group III. These PPIs were used as the positive samples for the training, validation, and test sets during model fine-tuning. To generate negative samples for each set, we also followed the previous procedure. Specifically, the negative samples were generated through random pairing protein monomers within each group after filtering out protein pairs sharing interacting partners, and a 10:1 negative-to-positive ratio was adopted for each set. For the HINT PPI datasets, where experimental monomer structures are not always available, we employed predicted monomer models from AlphaFoldDB as substitutes to generate the input features.

##### 3.3.2 Training protocol

The fine-tuning used the interaction labels to update model weights according to the interaction loss (Loss_interaction_), while preserving the model’s ability to predict inter-protein contacts by incorporating the contact maps of structure-informed PPIs as constraints using **μ** · **Loss**_**contact**_ with **μ** = **100**. Structure-informed PPIs were not used to train the interaction module during fine-tuning. Training followed the same protocol as in Section 3.2.2: PyTorch (v2.1.0) on a single NVIDIA A100 GPU, learning rate 1 × 10^-5^, AdamW optimizer, batch size 1, with learning rate reductions based on validation loss. The final model was selected based on the lowest validation loss. Notably, inter-protein contact maps were used only for training guidance and were not involved in model validation or selection.

### 4. Training of D-SCRIPT, Topsy-Turvy, TT3D, and TUnA

The training scripts for D-SCRIPT, Topsy-Turvy, and TT3D were downloaded from https://dscript.csail.mit.edu/. Both the initial training and the fine-tuning of these models were implemented under the default settings provided in the official tutorial (https://d-script.readthedocs.io/en/stable/usage.html#training) using the same datasets as the training of PLMDA-PPI. The training scripts for TUnA were downloaded from https://github.com/Wang-lab-UCSD/TUnA. The initial training and fine-tuning of the model were also implemented under the default settings using the same datasets as the training of PLMDA-PPI.

### 5. Implementation of AF2Complex, RF2-Lite, and RF2-PPI

AF2Complex was obtained from https://github.com/FreshAirTonight/af2complex and executed using the settings provided in example1.sh. RF2-Lite and RF2-PPI were obtained from https://github.com/SNU-CSSB/RF2-Lite and https://github.com/CongLabCode/RoseTTAFold2-PPI, respectively, and implemented under default settings using paired MSAs generated by ColabFold (https://github.com/sokrypton/ColabFold).

## Data Availability

The training and test data for this work can be accessed at https://github.com/ChengfeiYan/PLMDA-PPI/tree/main/data.

## Code Availability

The code for training and implementing PLMDA-PPI can be accessed at https://github.com/ChengfeiYan/PLMDA-PPI.

## Acknowledgements

The work was supported by the National Natural Science Foundation of China (grant number: 32571438 and 32101001) and the Major project of Guangzhou National Laboratory (grant number: GZNL2023A03007). Computational resources were provided by the HPC platform of Huazhong University of Science and Technology. The authors also utilized ChatGPT to assist in linguistic refinement and to enhance the clarity and readability of this manuscript.

## Author Contributions

C.Y. and S.-Y.H. conceived the project. C.Y. designed the experiments and supervised the study. S.D. and X.W. performed the experiments. S.D., M.Z., and C.Y. analyzed the data. S.D., X.W., and C.Y. drafted the manuscript. C.Y. and S.-Y.H. revised the manuscript. All authors discussed the results and reviewed the manuscript.

## Ethics Declarations

The authors declare no competing interests.

## Supplementary

**Figure S1.**
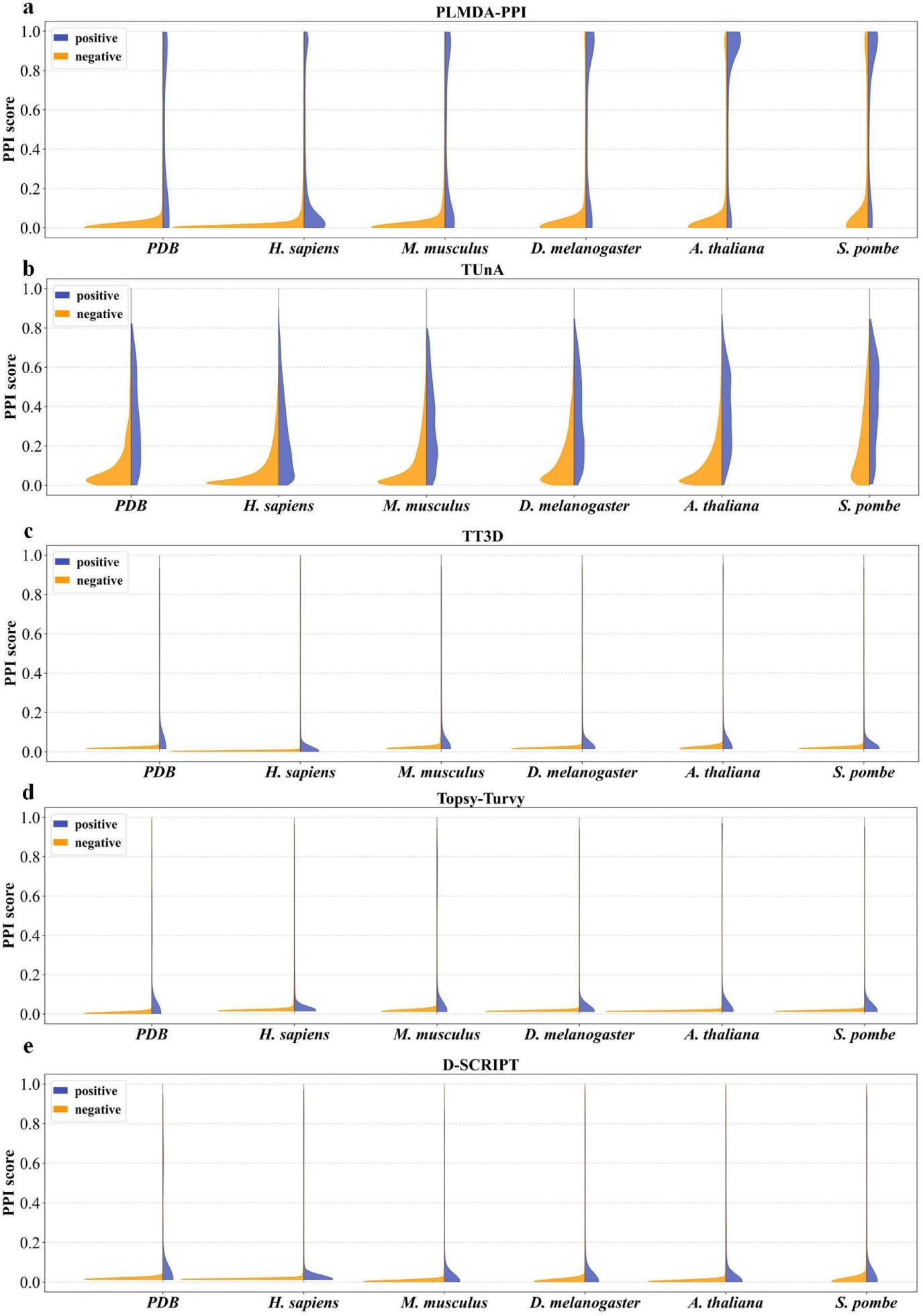
Distributions of PPI scores predicted by PLMDA-PPI (a), TUnA (b), TT3D (c), Topsy-Turvy (d), and D-SCRIPT (e) after initial training on structure-informed PPIs, for positive and negative samples in the validation set derived from structure-informed PPIs in PDB and the HINT test datasets across multiple species.

**Figure S2.**
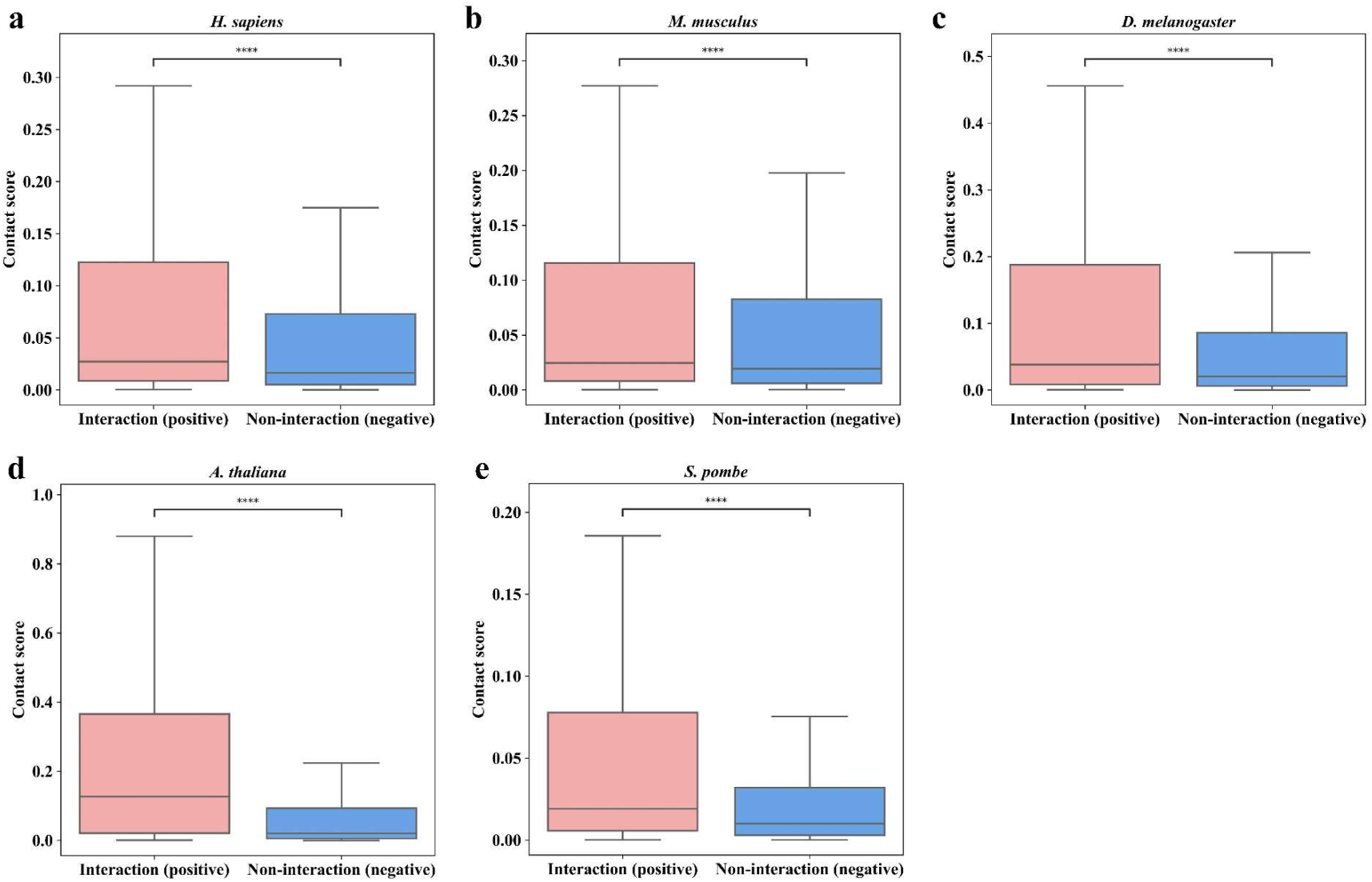
Comparison of the distributions of the mean contact score of the top 50 predicted residue pairs by PLMDA-PPI after initial training on structure-informed PPIs, for positive and negative samples in the HINT test datasets across multiple species. Asterisks indicate statistically significant differences based on the Mann-Whitney U test *p*-values: \*\*\*\**p*≤0.0001 (*H. sapiens*: *p*=5.8e-35, *M. musculus*: *p*=2.3e-12, *D. melanogaster: p*=5.9e-21, *A. thaliana*: *p*=7.8e-270, and *S. pombe*: *p*=6.1e-25).

**Figure S3.**
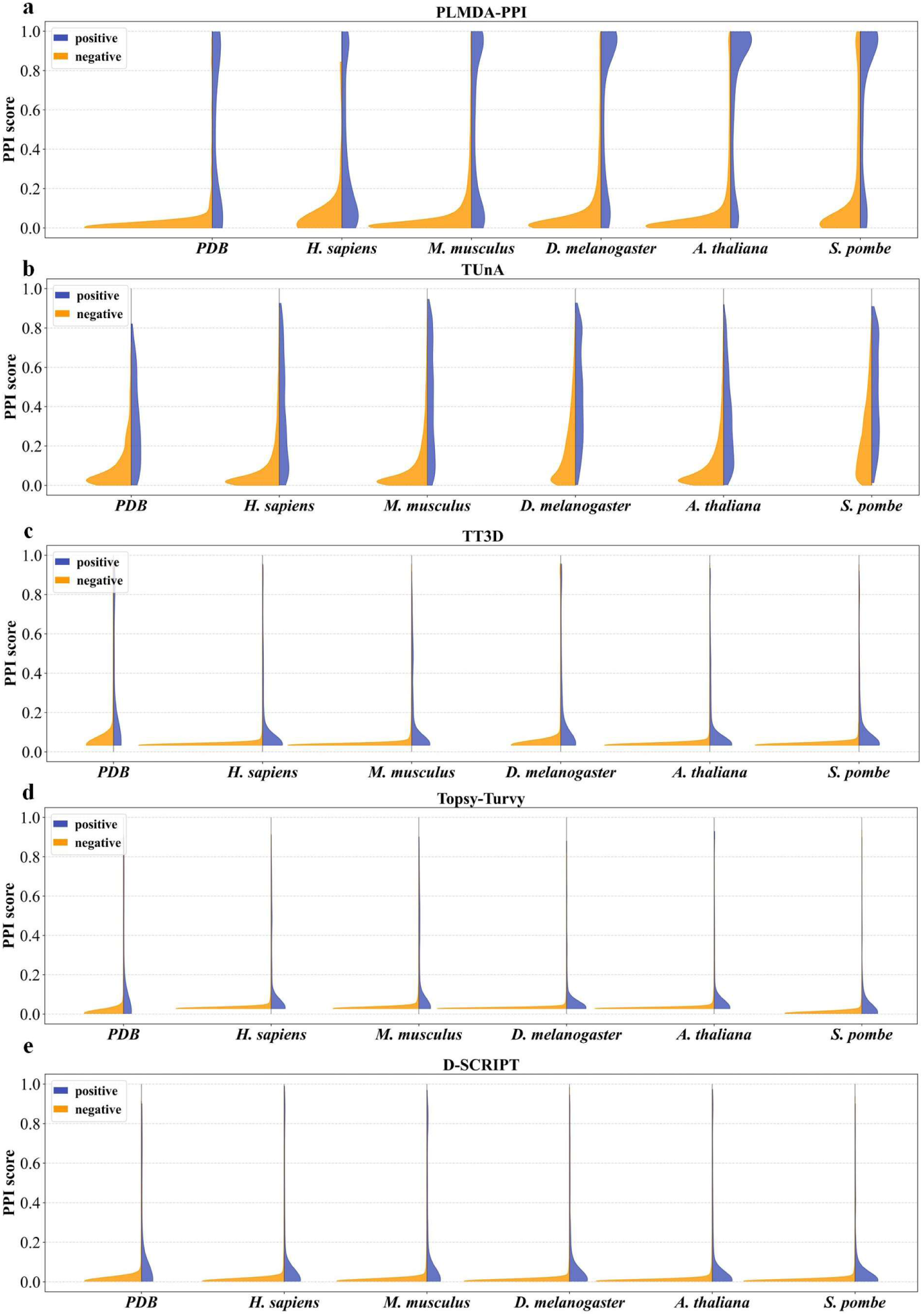
Distributions of PPI scores predicted by PLMDA-PPI (a), TUnA (b), TT3D (c), Topsy-Turvy (d), and D-SCRIPT (e) after transfer learning, for positive and negative samples in the validation set derived from structure-informed PPIs in PDB and the HINT test datasets across multiple species.

**Figure S4.**
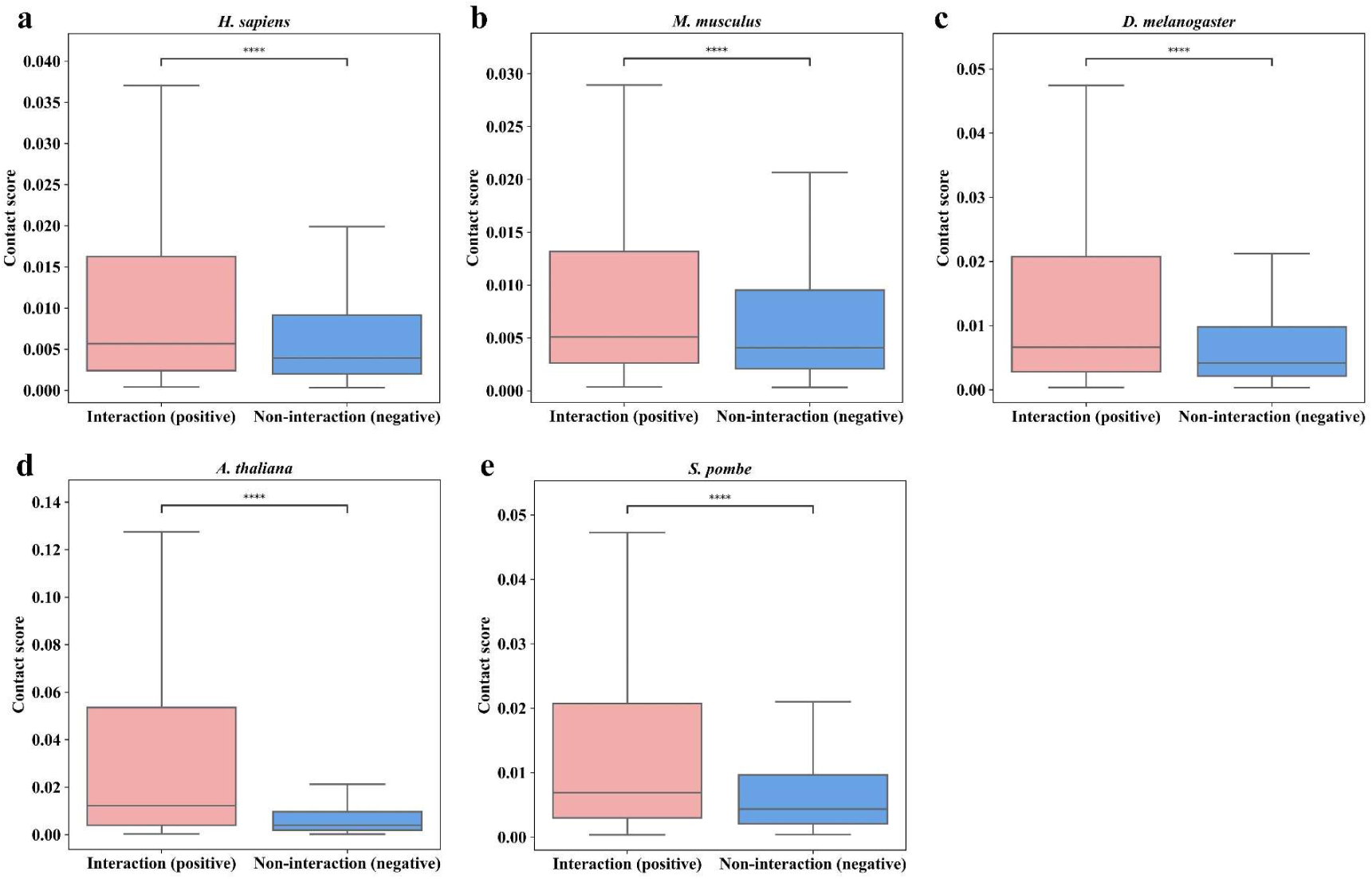
Comparison of the distributions of the mean contact score of the top 50 predicted residue pairs by PLMDA-PPI after transfer learning, for positive and negative samples in the HINT test datasets across multiple species. Asterisks indicate statistically significant differences based on the Mann-Whitney U test *p*-values: \*\*\*\**p*≤0.0001 (*H. sapiens*: *p*=2.1e-24, *M. musculus*: *p*=1.0e-25, *D. melanogaster*: *p*=2.2e-34, *A. thaliana*: *p*=1.9e-276 and *S. pombe*: *p*=1.4e-26).

**Figure S5.**
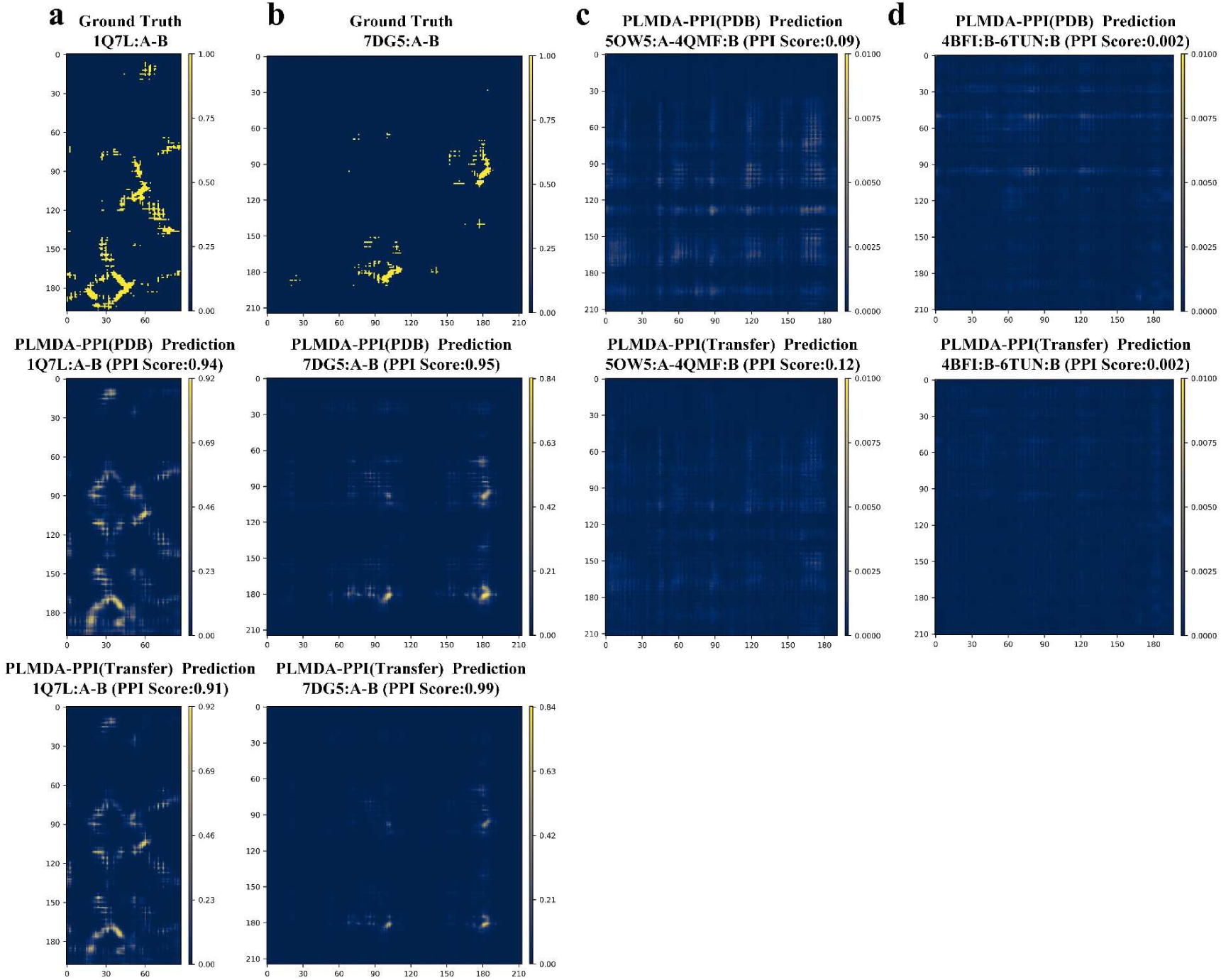
(a)∼(b) The experimental inter-protein contact maps (upper panels) and the inter-protein contact maps predicted by PLMDA-PPI before (middle panels) and after (lower panels) transfer learning for two positive samples ((a) PDB 1Q7L:A-B, (b) 7DG5:A-B). (c)∼(d) The inter-protein contact predicted by PLMDA-PPI before (upper panels) and after (lower panels) transfer learning for two negative samples ((c) PDB 5OW5:A-4QMF:B, (d) PDB 4BFI:B-6TUN:B)

**Figure S6.**
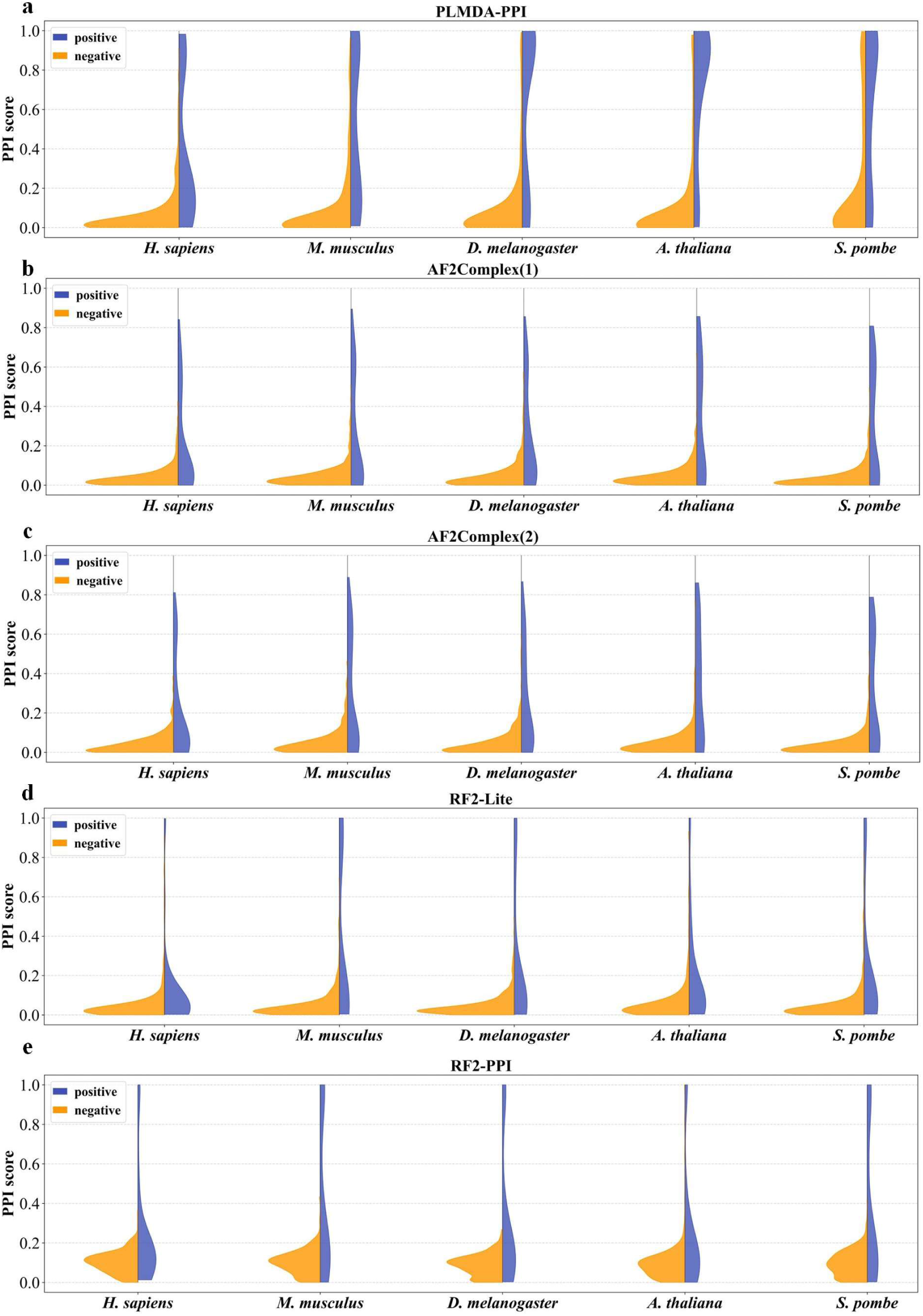
Distributions of PPI scores predicted by PLMDA-PPI (a), AF2Complex(1) (b), AF2Complex(2) (c), RF2-Lite (d), and RF2-PPI (e), for positive and negative samples sampled from original HINT test sets across multiple species.

**Table S1.**
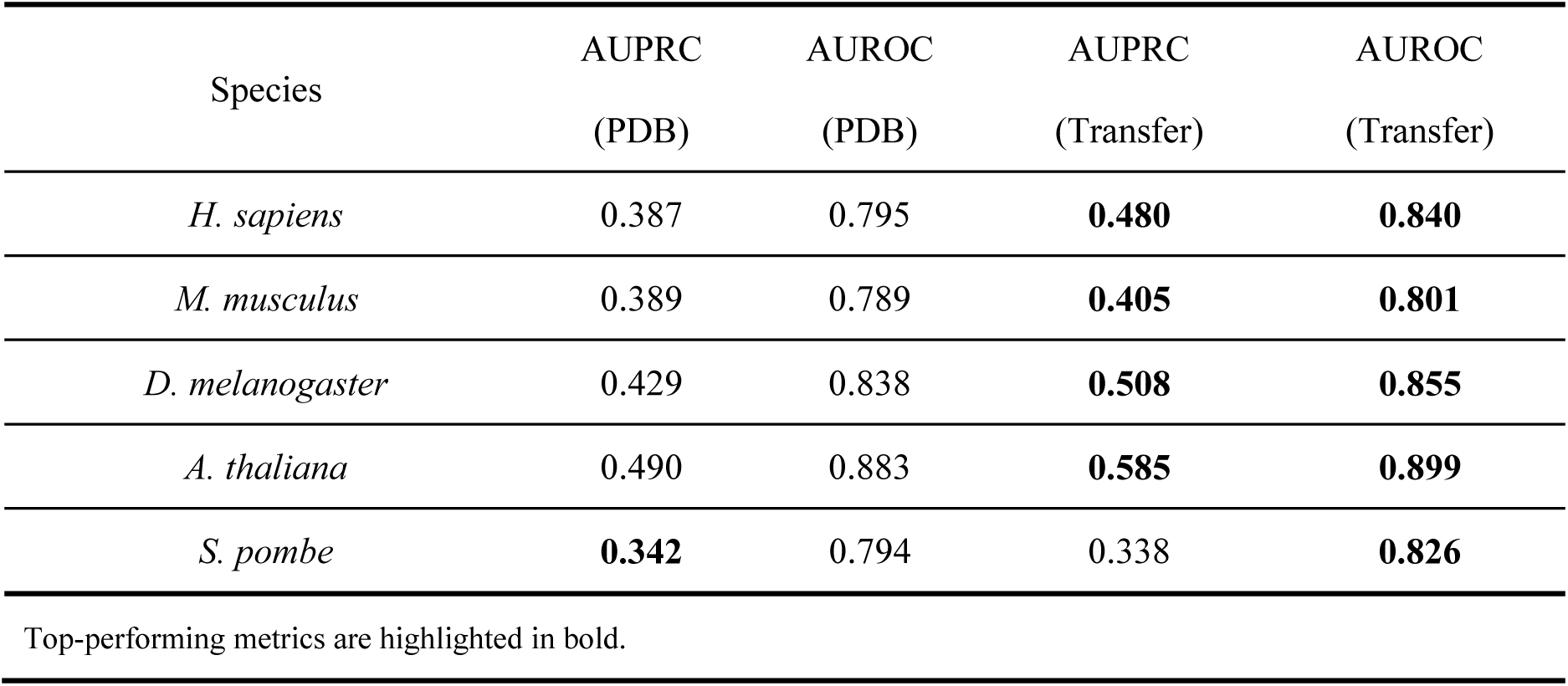
Performance comparison of PLMDA-PPI before and after HINT fine-tuning using HINT datasets across multiple species, following the exclusion of PPIs involving any protein sharing more than 40% sequence identity with proteins in the *H. sapiens* training set.

**Table S2.** Performances of the models when trained directly on the HINT *H. sapiens* PPI training set.

| Species | Model | AUPRC | AUROC |
| --- | --- | --- | --- |
| *H. sapiens* | D-SCRIPT | 0.197 | 0.74 |
|  | Topsy-Turvy | 0.212 | 0.709 |
|  | TT3D | 0.227 | 0.706 |
|  | TUnA | 0.313 | 0.791 |
|  | PLMDA-PPI | **0.400** | **0.812** |
| *M. musculus* | D-SCRIPT | 0.169 | 0.713 |
|  | Topsy-Turvy | 0.16 | 0.63 |
|  | TT3D | 0.196 | 0.688 |
|  | TUnA | 0.311 | 0.803 |
|  | PLMDA-PPI | **0.393** | **0.838** |
| *A. thaliana* | D-SCRIPT | 0.134 | 0.635 |
|  | Topsy-Turvy | 0.127 | 0.576 |
|  | TT3D | 0.153 | 0.593 |
|  | TUnA | 0.238 | 0.738 |
|  | PLMDA-PPI | **0.313** | **0.829** |
| *D. melanogaster* | D-SCRIPT | 0.123 | 0.615 |
|  | Topsy-Turvy | 0.141 | 0.563 |
|  | TT3D | 0.239 | 0.685 |
|  | TUnA | 0.210 | 0.748 |
|  | PLMDA-PPI | **0.378** | **0.828** |
| *S. pombe* | D-SCRIPT | 0.119 | 0.597 |
|  | Topsy-Turvy | 0.141 | 0.573 |
|  | TT3D | 0.204 | 0.66 |
|  | TUnA | 0.238 | 0.684 |
|  | PLMDA-PPI | **0.377** | **0.770** |

Top-performing metrics are highlighted in bold.

## References

1. Stelzl, U. et al. A human protein-protein interaction network: A resource for annotating the proteome. Cell 122, 975–68 (2005).

2. Huttlin, E.L. et al. The BioPlex Network: A Systematic Exploration of the Human Interactome. Cell 162, 425–40 (2015).

3. Mintseris, J. & Weng, Z. Structure, function, and evolution of transient and obligate protein-protein interactions. Proc. Natl. Acad. Sci. U. S. A. 102, 10930–5 (2005).

4. Drew, K., Wallingford, J.B. & Marcotte, E.M. hu.MAP 2.0: integration of over 15,000 proteomic experiments builds a global compendium of human multiprotein assemblies. Mol. Syst. Biol. 17, e10016 (2021).

5. Burke, D.F. et al. Towards a structurally resolved human protein interaction network. Nat. Struct. Mol. Biol. 30, 216–5 (2023).

6. Vidal, M. How much of the human protein interactome remains to be mapped? Science Signaling 9, eg7 (2016).

7. Luck, K. et al. A reference map of the human binary protein interactome. Nature 580, 402–8 (2020).

8. Snider, J. et al. Fundamentals of protein interaction network mapping. Mol. Syst. Biol. 11, 848 (2015).

9. Petrey, D., Zhao, H., Trudeau, S.J., Murray, D. & Honig, B. PrePPI: A Structure Informed Proteome-wide Database of Protein-Protein Interactions. J. Mol. Biol. 435, 168052 (2023).

10. Zhang, J., Durham, J. & Qian, C. Revolutionizing protein-protein interaction prediction with deep learning. Curr. Opin. Struct. Biol. 85, 102775 (2024).

11. Humphreys, I.R. et al. Computed structures of core eukaryotic protein complexes. Science 374, eabm4805 (2021).

12. Hu, L., Wang, X., Huang, Y., Hu, P. & You, Z. A survey on computational models for predicting protein-protein interactions. Brief. Bioinform. 22, bbab036 (2021).

13. Szklarczyk, D. et al. The STRING database in 2021: customizable protein-protein networks, and functional characterization of user-uploaded gene/measurement sets. Nucleic Acids Res. 49, D605–12 (2021).

14. Bernett, J., Blumenthal, D.B. & List, M. Cracking the black box of deep sequence-based protein-protein interaction prediction. Brief. Bioinform. 25, bbae076 (2024).

15. Park, Y. & Marcotte, E.M. Flaws in evaluation schemes for pair-input computational predictions. Nat. Methods 9, 1134–6 (2012).

16. Lannelongue, L. & Inouye, M. Pitfalls of machine learning models for protein-protein interaction networks. Bioinformatics 40, btae012 (2024).

17. Li, S. et al. MONN: A Multi-objective Neural Network for Predicting Compound-Protein Interactions and Affinities. Cell Syst. 10, 308–22(2020).

18. Nam, H., Ha, J. W. & Kim, J. Dual attention networks for multimodal reasoning and matching. in Proceedings - 30th IEEE Conference on Computer Vision and Pattern Recognition, CVPR 2017 vols 2017-January (2017).

19. Si, Y. & Yan, C. Protein language model-embedded geometric graphs power inter-protein contact prediction. eLife 12, RP92184 (2023).

20. Berman, H.M. et al. The Protein Data Bank. Nucleic Acids Res. 28, 235–242 (2000).

21. Das, J. & Yu, H. HINT: High-quality protein interactomes and their applications in understanding human disease. BMC Syst. Biol. 6, 1–12 (2012).

22. Sledzieski, S., Singh, R., Cowen, L. & Berger, B. D-SCRIPT translates genome to phenome with sequence-based, structure-aware, genome-scale predictions of protein-protein interactions. Cell Syst. 12, 969–82 (2021).

23. Singh, R., Devkota, K., Sledzieski, S., Berger, B. & Cowen, L. Topsy-Turvy: integrating a global view into sequence-based PPI prediction. Bioinformatics 38, i264–72 (2022).

24. Sledzieski, S., Devkota, K., Singh, R., Cowen, L. & Berger, B. TT3D: Leveraging precomputed protein 3D sequence models to predict protein - protein interactions. Bioinformatics 39, btad663 (2023).

25. Ko, Y.S., Parkinson, J., Liu, C. & Wang, W. TUnA: an uncertainty-aware transformer model for sequence-based protein-protein interaction prediction. Brief. Bioinform. 25, bbae359 (2024).

26. Gao, M. & Skolnick, J. Improved deep learning prediction of antigen-antibody interactions. Proc. Natl. Acad. Sci. 121, e2410529121 (2024).

27. Gao, M., Nakajima An, D., Parks, J.M. & Skolnick, J. AF2Complex predicts direct physical interactions in multimeric proteins with deep learning. Nat. Commun. 13, 1744 (2022).

28. Humphreys, I.R. et al. Protein interactions in human pathogens revealed through deep learning. Nat. Microbiol. 9, 2642–2652 (2024).

29. Zhang, J. et al. Predicting protein-protein interactions in the human proteome. Science 390, eadt1630 (2025).

30. Jumper, J. et al. Highly accurate protein structure prediction with AlphaFold. Nature 596, 583–9 (2021).

31. Evans, R. et al. Protein complex prediction with AlphaFold-Multimer. bioRxiv 2021.10.04.463034 (2021).

32. Efficient and accurate prediction of protein structure using RoseTTAFold2. bioRxiv 2023.05.24.542179 (2023).

33. Si, Y. & Yan, C. Improved protein contact prediction using dimensional hybrid residual networks and singularity enhanced loss function. Brief. Bioinform. 22, bbad039 (2021).

34. Si, Y. & Yan, C. Improved inter-protein contact prediction using dimensional hybrid residual networks and protein language models. Brief. Bioinform. 24, bbad039 (2023).

35. Rives, A. et al. Biological structure and function emerge from scaling unsupervised learning to 250 million protein sequences. Proc. Natl. Acad. Sci. U. S. A. 118, e2016239118 (2021).

36. Varadi, M. et al. AlphaFold Protein Structure Database: massively expanding the structural coverage of protein-sequence space with high-accuracy models. Nucleic Acids Res. 50, D439–44 (2022).

37. van Kempen, M. et al. Fast and accurate protein structure search with Foldseek. Nat. Biotechnol. 42, 243–6 (2024).

38. Saito, T. & Rehmsmeier, M. The Precision-Recall Plot Is More Informative than the ROC Plot When Evaluating Binary Classifiers on Imbalanced Datasets. PLoS One 10, e0118432 (2015).

39. Mirdita, M. et al. ColabFold: making protein folding accessible to all. Nat. Methods 19, 679–682 (2022).

40. Li, W. & Godzik, A. Cd-hit: a fast program for clustering and comparing large sets of protein or nucleotide sequences. Bioinformatics 22, 1658–9 (2006).

41. Hsu, C. et al. Learning inverse folding from millions of predicted structures. Proceedings of the 39th International Conference on Machine Learning, PMLR 162:8946-8970 (2022).

42. Johnson, L.S., Eddy, S.R. & Portugaly, E. Hidden Markov model speed heuristic and iterative HMM search procedure. BMC Bioinformatics 11, 1–8 (2010).

43. Suzek, B.E., Huang, H., McGarvey, P., Mazumder, R. & Wu, C.H. UniRef: comprehensive and non-redundant UniProt reference clusters. Bioinformatics 23, 1282–1288 (2007).

44. Rao, R. et al. MSA Transformer. Proceedings of the 38th International Conference on Machine Learning, PMLR 139:8844–8856 (2021).

45. Jing, B., Eismann, S., Suriana, P., Townshend, R. J. L. & Dror, R. O. Learning from Protein Structure with Geometric Vector Perceptrons. in ICLR 2021 - 9th International Conference on Learning Representations (2021).

